# A role for the V0 sector of the V-ATPase in neuroexocytosis: exogenous V0d blocks complexin and SNARE interactions with V0c

**DOI:** 10.1101/2023.01.31.526435

**Authors:** Christian Leveque, Yves Maulet, Qili Wang, Marion Rame, Sumiko Mochida, Marion Sangiardi, Youssouf Fahamoe, Cécile Iborra, Michael Seagar, Nicolas Vitale, Oussama El Far

## Abstract

V-ATPase is an important factor in synaptic vesicle acidification and is implicated in synaptic transmission. Rotation of the extra-membranous V1 sector drives proton transfer through the membrane-embedded multi-subunit V0 sector of the V-ATPase. Intra-vesicular protons are then used to drive neurotransmitter uptake by synaptic vesicles. V0a and V0c, two membrane subunits of the V0 sector have been shown to interact with SNARE proteins and their photo-inactivation rapidly impairs synaptic transmission. V0d, a soluble subunit of the V0 sector strongly interacts with its membrane embedded subunits and is crucial for the canonic proton transfer activity of the V-ATPase. Our investigations show that the loop 1.2 of V0c interacts with complexin, a major partner of the SNARE machinery and that V0d1 binding to V0c inhibits this interaction, as well as V0c association with SNARE complex. Injection of recombinant V0d1 in rat superior cervical ganglion neurons rapidly reduced neurotransmission. In chromaffin cells, V0d1 overexpression and V0c silencing modified in a comparable manner several parameters of unitary exocytotic events. Our data suggest that V0c subunit promotes exocytosis via interactions with complexin and SNAREs and that this activity can be antagonized by exogenous V0d.

## Introduction

V-ATPase is an important player in acidifying intracellular compartments in eukaryotes. The molecular integrity of this enzyme guarantees energy-dependent proton transfer into specific compartments. Many intracellular organelles such as endosomes, trans-Golgi, secretory granules, synaptic vesicles, and lysosomes are all critically dependent, for their function, on an acidic pH generated by the V-ATPase [1]. This pH sensor [2] and mechano-chemical energy transducer [3] is composed of two reversibly attached intertwined V1 and V0 domains, each having a complex subunit composition [4] [5]. The structural organization of this enzyme is intricate and the reversible association of V0 and V1 regulates the coupling of ATPase activity to proton transport and consequent acidification of membrane compartments [6] [7]. Synaptic vesicle acidification is functionally associated with vesicular neurotransmitter uptake and it has been reported that the extra-membranous V1 dissociates from fully loaded vesicles Synaptic vesicles acidification is believed to induce the dissociation of the extra-membranous V1 sector [7]. The V1 domain performs the ATPase activity while proton transport takes place through the membrane-embedded V0 domain. The latter is composed of a tight assembly of several subunits (V0a, V0d, V0e, Ac45 and ATP6AP2), all surrounding a very hydrophobic rotor composed of several copies of V0c and a single V0c” subunit. V0e, Ac45 and ATP6AP2 are single pass transmembrane proteins, V0c and c” are tetraspans with two cytosolic loops and V0a possesses 8 TMD and a prominent cytosolic N-terminus. Although the boxing-glove-shaped V0d [8] has no transmembrane domain (TMD) [9] [5], it is a stable component of the V0 domain [10] [11] and is required for assembly of V-ATPase complex [12]. Yeast V0d participates in the so called “stator” [13] and has been shown to interact with the large N-terminal domain of V0a [14] [15] [16] following V1 dissociation, which results in an auto-inhibited form of V0 and that is no longer capable of proton translocation. In humans, there are two isoforms of V0d: a ubiquitous V0d1 and second isoform, V0d2, selectively expressed in osteoclasts, lung, kidney and epididymis [17]. Both isoforms interact with the central stalk V1 subunits V1D and V1F as well as with the V0c rotor [9] [5] [18]. V0d1 knockout in the mouse is embryonic lethal, highlighting its important physiological role [19]. Consequently, it has been proposed that the V0d subunit could couple the cytosolic part of the enzyme to its membrane embedded component and therefore potentially couple ATP hydrolysis to proton transport [20].

Independently of its critical role in proton translocation, V0 domain is implicated in exocytosis [21], neurotransmitter release [22] [23], and also in numerous membrane fusion events in intracellular compartments [24] [25] [26] [27]. However, genetic inactivation studies cannot be used to examine the role of the V0 domain in membrane fusion events since proton translocation is also needed for vesicle loading with neurotransmitters and hormones. At the molecular level, neurotransmitter release requires the formation of the soluble N-ethylmaleimide-sensitive factor attachment protein receptors (SNARE) complex composed of two proteins of the presynaptic plasma membrane, syntaxin and SNAP-25, and a synaptic vesicle membrane protein, VAMP / synaptobrevin [28]. This minimal fusion machinery is regulated by other factors such as the small soluble presynaptic protein complexin that binds assembled SNARE complexes and may promote oligomerization of the SNARE complexes and fusion [29] [30]. Several studies have shown that V0 interacts with and potentially regulates SNARE proteins [31] [24] [32] [22] [33]. Inhibition of the V0c loop 3.4 interaction with VAMP2 was shown to significantly decrease neurotransmission and thus highlighted the importance of V0c interaction in modulating SNARE-dependent neurotransmission [33]. Chromophore-assisted light inactivation (CALI) later demonstrated that the photo-inactivation of the V0a subunit rapidly impaired neuronal synaptic transmission and catecholamine release from chromaffin cells [7]. In contrast, photo-inactivation of the V1 catalytic subunit A, very much like pharmacological inhibition of proton transport, induced a delayed inhibition of neurosecretion, strongly arguing that V0 regulates exocytosis independently from proton transport [7] without being directly involved in forming a membrane fusion pore [7] [34]. More recently, we showed, using CALI experiments, that photo-inactivation of V0c in CA3 pyramidal neurons, rapidly inhibits neurotransmitter release downstream of synaptic vesicle acidification, thus corroborating the importance of V0 domain subunits in directly modulating neurotransmission [35]. However, the exact function of the V-ATPase in regulating membrane fusion events remains a matter of debate.

In this study we discovered that V0c interacts directly with complexin as well as with a mature assembly containing complexin and the trimeric SNARE complex. This interaction is mediated by the first cytosolic loop 1.2 of V0c that also mediates V0c interaction with V0d1. We show that V0c bound to V0d is no longer available to interact with complexin and the SNARE complex. Similar to V0c silencing, V0d1 overexpression or its intraneuronal injection inhibited exocytosis. Altogether these results bring new insight for the role of V-ATPase in exocytosis and reinforce the hypothesis that the V0 sector of the V-ATPase is an important modulator of SNARE-dependent neurosecretion.

## Materials and Methods

### Reagents

Unless otherwise stated, chemicals were principally from Sigma-Aldrich®. Oligonucleotides were from Eurofins genomics, Glutathione Sepharose and CM5 sensor chips for Surface Plasmon Resonance (SPR) were from Cytiva. Protease inhibitors were either from GE Healthcare (cOmplete) or Thermo Scientific (Halt− protease inhibitor cocktail). Lipids were from Aventi Polar. Polyclonal anti V0d1 was from Proteintech and anti actin was from Sigma-Aldrich. Anti-HSV rabbit polyclonal antibody was from Abcam (ab19355), anti-T7 from Novagen and anti-GST polyclonal antibody was from GE-Healthcare. Anti-syntaxin 1 (10H5) [36], anti-complexin 1 (SP33) and 2 (LP27) [37] [38] and anti-SNAP-25 BR05 [39] antibodies were a generous gift from M. Takahashi. Monoclonal anti-rat/mouse VAMP2 (6F9) (aa 2-SATAATVPPAAPAGEGG -18) and anti-rat/mouse SNAP-25 (6C11) (aa 196-NQRATKMLGSG -206) were produced and protein-A purified by Genecust. V0c L1.2 (aa 35-KSGTGIAAMSVMRPELIMKS -54) and L3.4 (aa 117-GVRGTAQQPRLF -155) peptides were synthesized by Genecust. Fos-Choline-12 (FC12) was purchased from Anatrace and CHAPS from Euromedex. Maxisorp ELISA plates were from Nunc. MEM, horse and fetal calf serum as well as Glutamax and penicillin streptomycin were from (Life Technologies™). Coverslips were from Amilabo. Petri dishes from Falcon ®, poly-D-lysine was from Sigma-Aldrich®. Neurosensor 510 (7-(Diethylamino)-4-(4-methoxyphenyl)-2-oxo-2H-1-benzopyran-3-carboxaldehyde) was purchased from (Tocris).

### Expression plasmids and cloning procedures

pET21-Complexin1-His plasmid construct was generated by cloning the rat complexin1 coding sequence in pET21 between Nde1 and Hind III restriction sites. Complexin1-GST expressing plasmid was constructed as follows: The Nco1 restriction site in pET28a was replaced by the one of Nde1 and the GST coding sequence was introduced downstream of the Not1 restriction site of this plasmid. Rat complexin 1 coding sequence was then inserted (Nde1-EcoR1) upstream of GST in this modified vector. Bacterial expression plasmids pET16-VampΔTM-Myc-His, pRSF-StxΔTM and pQE30-His-SNAP25 producing soluble SNARE proteins His6-tagged Vamp(1-96), untagged syntaxin 1 (1-265) and His6-tagged SNAP25 were described previously [40] [41]. His6-T7-HSV-tagged full length Plasmid pRSF-Duet-SNAP25-StxΔTM co-expressing untagged SNAP25 and syntaxin 1 (1-265) was obtained by successively inserting syntaxin (1-265) sequence into NdeI / XhoI sites of pRSF-Duet-1 (Novagen) and SNAP25 sequence into the NcoI / EcoRI sites of the resulting plasmid. His6-T7-HSV tagged full length V0c-subunit construct was previously described [33]. Full length V0c-L1.2s mutant was constructed using two overlapping PCR fragments. Both fragments overlapped in the loop 1.2 (L1.2) region and encoded the following scrambled L1.2 sequence (L1.2s): SMGITLGEIPARAMMVS. To amplify the 5’ half of the sequence, T7 was used as a forward primer and 5’-cgagaccat catggcccttgctgggatctcgcccagagtgatgcccatactCTTGGCTGTGCCATAGGC-3’ as reverse. The 3’ half was amplified using 5’-agtatgggcatcactctgggcgagatcccagcaagggccatgatggtctcgAAGTCCATCATCCCAGTGG as forward and 5’-gcgtcgacCTACTTTGTGGAGAGGATTAG-3’ as reverse. These two fragments were then mixed in the absence of any primer and the full length V0c L-1.2s was PCR amplified and cloned in pET28 using EcoR1 and Sal1 sites. The same procedure was used to generate full length V0c-L3.4s mutant. Both fragments overlapped in the loop 3.4 region and encoded the following scrambled L3.4 sequence (L3.4s): GQATVQPLGRRF. To amplify the 5’ half of the sequence, T7 was used as forward primer and 5’-gaatcgccggcccagaggctggacagtggcctgaccAGC ATCTCCGACAATGCC-3’ as reverse. The 3’ half was amplified using 5’-ggtcaggccactgtccagcctct gggccggcgattcGTGGGCATGATCCTGATCC-3’ as forward and 5’-gcgtcgacCTACTTTGTGGAGAGGATTAG-3’ as reverse. These two fragments were then mixed in the absence of any primer and the full length V0c-L-3.4s was PCR amplified and cloned in pET28 using EcoR1 and Sal1 sites. Rat V0d1 was amplified by PCR using a commercial Y2H adult rat brain plasmid cDNA library (Origine). EcoR1 and Sal1 sites were used for insertion in plasmid constructs (Forward: GCGAATTCTCGTTCTTCCCGGAGCTTT, reverse: GCGTCGACCTAGAAGATGGGGATAT AGTTG). Construction of GST-V0d1 expression plasmid was obtained by insertion of amplified V0d1 into EcoR1/Sal1 digested pGEX-5x-1. Construction of 6His-HA-V0d1 expression plasmid was performed as follows. HA tag sequence (YPYDVPDYA) was introduced by linker insertion at the 5’ side of pET28a (Novagen) MCS to generate pET28-HA-Nter. Forward (5’-p-GATCCTATCCTTATGATGTTCCTGATTATGCAG) and reverse primers (5’-p-AATTCTG CATAATCAGGAACATCATAAGGATAG) were annealed and ligated to BamH1/EcoR1 digested pET28a. EcoR1 / Sal1 sites were used to insert amplified full length V0d1. GST-HSV expression plasmid was constructed using a phosphorylated linker encoding the HSV tag (QPELAPEDPED) sequence and ligated to BamH1 / EcoR1 digested pGEX-4T1 MCS. Forward (5’-p-GATCCCAGCCTGAACTCGCTCCAGAAGACCCGGAAGATG) and reverse (5’-p-AATTCATCTTCCGGGTCTTCTGGAGCGAGTTCAGGCTGG) primers. Bicistronic pIRES-2-EGFP plasmid co-expressing myc-V0d1 was constructed using Xho1-Sal1 restriction sites. In the myc-V0d1-expressing pIRES-2-turbo-RFP, the coding sequence of EGFP was replaced by the one of turboRFP amplified by PCR from pINDUCER11 [42].

### Recombinant protein expression

Complexin1-His, complexin1-GST, His-Vamp(1-96) and t-SNAREs (syntaxin 1 (1-265) + His-SNAP25) were expressed in BL21 and purified as previously described [33,40]. Soluble trimeric SNARE complexes were produced by co-transfecting BL21 with pET16-VampΔTM-myc-His and pRSF-Duet-SNAP25-StxΔTM. Bacteria were grown in TB medium supplemented with both ampicillin and kanamycin. Expression was induced 4 hrs at 37°C by 0.5 mM isopropyl-thio-βD-galactoside. BL21 expressing V0d1 constructs were cultured in TB and protein expression was induced with 0.3 mM IPTG for 17 hours at 18°C. pET-28 containing 6His-V0c construct was transfected in the OverExpress− C43(DE3) bacterial strain (Avidis, France) and expression induced as for V0d1. All bacterial pellets were stored at -20°C before protein purification.

### Protein purification

Protease inhibitors were present in all homogenisation steps. Recombinant soluble trimeric SNARE complexes were obtained from 2l of induced culture. Purification was performed at room temperature when not otherwise stated. Pelleted bacteria (7 g) were resuspended in 25 ml of buffer H (sodium phosphate 50 mM, 0.5 M NaCl, 20 mM imidazole, pH 8) and lysed in a French press. The homogenate was centrifuged (200.000 g, 30 min, 4°C) and the supernatant incubated 1 hr at 4°C with 1.5 ml packed Ni-NTA Agarose beads (Qiagen). Beads were washed with buffer H adjusted to 50 mM imidazole. Purified proteins were eluted in 0.5 ml fractions by increasing imidazole concentration to 250 mM. Peak fractions were dialyzed against 20 mM Tris-HCl, 1.0 mM EDTA pH7.4 and loaded on a 1 ml HiTrap-Q column equilibrated in the same buffer on an AKTA-purifier system. Proteins were eluted with a 10 - 500 mM NaCl gradient in the dialysis buffer. The main peak fractions eluting at 375.0 mM NaCl were pooled. Proteins were quantified by Bradford assay and analysed on SDS-PAGE gel and western blot. Aliquots were stored at -20 °C. Bacterial pellets from 6His-V0c expression, were resuspended in wash buffer (50 mM Tris-HCl, pH 8.0, 1 mM EDTA) supplemented with 0.2 mg/ml lysozyme and subjected to French press. Homogenates were centrifuged 5 min at 2500 x g and the membrane fraction was isolated from supernatant by centrifuging at 200.000 x g for 37 min. Membrane were washed once by resuspension of the pellet in wash buffer and centrifugation. Membranes (5 mg/ml protein) were solubilized in 50 mM Tris-HCl, pH 8.0, 10 mM β-mercapto-ethanol, 2 % FosCholine-12 at 4°C for 1 hr. The buffer was then adjusted to 0.5 M NaCl, 20 mM imidazole and the recombinant V0c was purified over Ni-NTA beads (QIAGEN).

Bacterial pellets expressing V0d1 constructs were resuspended in 50 mM Tris-HCl pH 8.0, 150 mM NaCl and subjected to French Press. Insoluble material was eliminated by centrifugation 37 min at 200.000 x g. GST-V0d1 was purified by batch incubation of supernatant (1hr 4°C) with 1 ml Glutathione-Sepharose. The suspension was loaded on a disposable 10 ml column. Washing (20 mM HEPES pH7.4, 150mM NaCl, 0.1% Triton X-100 then 20 mM HEPES pH7.4, 150 mM NaCl) and elution (20 mM HEPES pH 7.4, 150 mM NaCl, 10 mM reduced glutathione) were performed manually.

For purification of 6His-HA-V0d1, the high-speed supernatant was adjusted to 0.5 M NaCl and 20 mM imidazole and batch incubated with 2 ml Ni-NTA-Sepharose beads (1hr, room temperature). The suspension was loaded on a disposable column, washed with 20 mM HEPES, 150 mM NaCl, in the presence of 20 mM and 40 mM imidazole successively and eluted by fractions in the presence of imidazole 0.5 M.

For both GST- and 6His-V0d1 preparations, protein-containing fractions were pooled, desalted against 10 mM HEPES pH 7.4 on Zeba-spin columns (Thermo scientific, USA), aliquoted, fast frozen in liquid N_2_ and stored in aliquots at -80°C. Protein concentrations were determined by parallel Coomassie staining and A_280_ absorbance. The very high hydrophobicity of V0c renders less accurate its protein assay by classical methods. Therefore, all 6His-HSV-V0c protein preparations were submitted to comparative relative quantification on Western blot using anti HSV antibodies and GST-HSV fusion protein as a standard.

### Preparation of rat brain extracts

Rat brain fraction enriched in plasma membrane (LP1) was purified as described [43]. Briefly, proteins were solubilized at 1.5 mg/ml in 25 mM Tris pH7.4, 150 mM NaCl containing 1.5 % CHAPS, 5 mM EDTA and proteases inhibitors before centrifugation 1 h at 14000 x g and filtration through 0.22 µm filters.

### V-ATPase mediated proton uptake

Proton uptake experiments were performed as previously described [33]. Briefly, rat brains were homogenized in 0.32 M sucrose, 10 mM HEPES pH 7.4, 0.2 mM EGTA in the presence of protease inhibitor cocktail and the post-nuclear supernatant was layered onto a 0.8 M sucrose cushion and centrifuged 20 min at 257.000 x g. Synaptosomal pellet was resuspended in hypotonic buffer (10 mM Tris-HCl pH 8.5, 1 mM PMSF) and centrifuged 20 min at 40.000 x g. The synaptic vesicle-enriched supernatant was adjusted to 10 mM Tris-HCl pH 8.5, 60 mM sucrose, 140 mM KCl, 2 mM MgCl_2_, 50 μM EGTA (assay buffer).

Synaptic vesicles were preincubated with 2 μM acridine orange (AO) before triggering proton uptake by the addition of 500 μM Mg-ATP. ATP-dependent proton transport was monitored, using a Biotek Sirius HT injector plate reader, by the quenching of AO fluorescence at 525 nm with data acquisition every 15 seconds.

### ELISA

HSV-V0c was probed with anti-HSV antibody (1/4000), GST-V0d1 and GST with goat anti-GST polyclonal antibody (1/4000). VAMP2, syntaxin and complexin were respectively probed with monoclonal antibodies 6F9 [33], 10H5 and SP33 at 1.0-1.2 µg/ml. Proteins were immobilized overnight at 4°C on Maxisorb™ (Nunc) 96 wells microplates at 1µg/well in 100 µl NaHCO_3_ 0.1 M pH 8.9. After a 1 hr blocking step at 37°C in 200 µl binding buffer (BB: 50 mM Tris-HCl pH 7.5, 150 mM NaCl, 3% BSA w/v), incubations with interacting proteins (100 µl) at the indicated concentrations were performed overnight at 4°C in BB supplemented with 0.1% Triton X-100. For sequential binding, incubation with the first interactant was performed for 6 h at 4°C, then the solution was replaced by a fresh one containing both the first and the second interactant (V0c or V0d) and the plates were further incubated overnight at 4°C. After interactions, plates were washed three times with 200 µl/well of washing buffer (WB: 50 mM Tris-HCl pH 7.5, 150 mM NaCl, 0.1% BSA w/v, 0.1%Triton X-100) and incubated 1hr at 4°C with 100 µl of specific primary antibodies in BB supplemented with 0.1% Triton X-100. Wells were washed (3 × 200 µl/well WB) and incubated 1 hr at 4°C with 100 µl horse radish peroxidase-coupled anti-IgG (either donkey anti-rabbit, goat anti-Mouse or rabbit anti-goat, depending on the primary antibody). After three washes, plates were revealed with TMB solution standard (Uptima, Interchim, France) according to supplier specifications. Colour development was stopped with 100 µl of 1.0 M H_2_SO_4_ and absorbance read at 450 nm. Measurements were systematically performed in triplicates. Unless otherwise stated, blanks were always subtracted in the represented histograms. GST was used as a control binding partner when GST-V0d1 binding was measured. Signal in GST wells was either subtracted from GST-V0d1 signal or shown in parallel.

### Surface plasmon resonance analysis

SPR experiments were performed at 25°C using Biacore 3000 (GE healthcare) or Biacore T200 (Cytiva). The running buffer was HBS (10 mM HEPES pH 7.4, 150 mM NaCl) supplemented or not with CHAPS 0.2%. Non-specific binding on control flow cells was automatically subtracted from experimental measurements to yield the specific signal. Proteins were coupled on sensor chips CM5 (Cytiva) or CMDP from Xantec (2D carboxylmethyldextran). About 25 fmoles of GST, Complexin1-GST or GST-V0d were covalently immobilized using amine coupling chemistry at pH 5. Analytes were injected at 10 µl/min and blank run injections of running buffer were performed in the same condition permitting to yield double-subtracted sensorgrams using Biaevaluation 4.2 software (GE Healthcare).

SPR detection of native SNAREs/complexin on immobilized recombinant V0c: Solubilized proteins from LP1 were diluted 5-fold in running buffer (25 mM Tris-HCl pH 7.4, 150 mM NaCl, 0.2% CHAPS) and injected over 2 flow cells (CMDP chip) in series including a control flow cell functionalized with irrelevant mouse antibodies and another one with anti HSV antibodies that previously captured HSV-V0c.

### Superior cervical ganglion neurons and synaptic transmission recordings

6-8 week cultures, EPSP recording and injection of recombinant His-V0d (6.5 µM) or BSA (20 µM) were performed as described previously [44]. EPSPs were recorded at 0.1 Hz. The peak amplitudes were normalized to the values before injection. The averaged and smoothed values (Origin 7.5) obtained from an eight-point moving average algorithm were plotted against recording time with t=0 corresponding to the start of 3 min presynaptic injection of His tagged V0d. Data are mean ± SEM and statistical significance was evaluated using a two-tailed Man Whitney U test.

### Chromaffin cell culture and catecholamine release recordings

Freshly dissected primary bovine chromaffin cells were cultured in DMEM in the presence of 10 % fetal calf serum, 10 µM cytosine arabinoside, 10 µM fluorodeoxyuridine and antibiotics as described previously [45]. Plasmids and siRNAs were introduced into chromaffin cells (5 × 10^6^ cells) by Amaxa Nucleofactor systems (Lonza) according to manufacturer’s instructions and as described previously [46]. 48h/96h after transfection, catecholamine secretion was evoked by applying K^+^ (100 mM) in Locke’s solution without ascorbic acid for 10 s to single cells by mean of a glass micropipette positioned at a distance of 30-50 μm from the cell. Electrochemical measurements of catecholamine secretion were performed using 5 µm diameter carbon-fiber electrodes (ALA Scientific) held at a potential of +650 mV compared with the reference electrode (Ag/AgCl) and approached closely to the transfected cells essentially as described previously [47]. Amperometric recordings were performed with an AMU130 (Radiometer Analytical) amplifier, sampled at 5 kHz, and digitally low-pass filtered at 1 kHz. Analysis of amperometric recordings was done as described previously [48], allowing automatic spike detection and extraction of spike parameters. The number of amperometric spikes was counted as the total number of spikes with an amplitude > 5 pA. Data were analyzed using SigmaPlot 13 software. In the figure legends, n represents the number of cells analyzed. Statistical significance has been assessed using t-test as data fulfilled requirements for parametric tests.

### Catecholamine content measurement

Chromaffin cells were incubated for 30 min with 0.5 µM of Neurosensor 510 (NS510) at 37°C and then washed to remove excess sensor before fixation. Labelled cells were visualized using a Leica SP5II confocal microscope and quantification of NS510 signal was performed in individual cells.

## Results

### V0c loop 1.2 binds complexin

We have previously shown that the cytosolic loop (L3,4) linking transmembrane regions 3 and 4 of V0c binds VAMP2 and that this interaction modulates neurotransmission [33]. We therefore screened for other potential V0c binding partners. Recombinant V0c was immobilized on the chip of an SPR apparatus and a detergent extract of a lysed rat brain synaptosomal fraction was injected into the flow cell (Supplemental Fig 1a). Bound proteins were then detected by injecting monoclonal antibodies against proteins involved in exocytosis (Supplemental Fig. 1b). In addition to the previously reported SNARE proteins (VAMP2, syntaxin 1, and SNAP25 [49] [33]), complexin1 and 2 were specifically identified in association with V0c (Supplemental Fig. 1b). As complexin could be indirectly associated with V0c due to its association with the SNARE complex [49], we used purified recombinant proteins to investigate whether a direct interaction occurred. Further investigations of complexin / V0c interactions were performed using complexin 1 hereafter referred to as complexin. When complexin was immobilized on ELISA plates, a strong binding of V0c (20 nM) was detected (Fig. 1a) while up to 1μM of V0c did not generate significant binding to BSA-blocked ELISA wells in the absence of complexin (Fig. 1a and Supplemental Fig. 2). V0c binding to complexin was not inhibited in the presence of VAMP2 suggesting that VAMP2 and complexin interact with V0c through distinct domains (Fig. 1a). In order to identify the molecular determinants on V0c that are important for complexin binding, we probed the implication of both cytosolic V0c loops that link respectively V0c TMDs 1 & 2 and 3 & 4. For this purpose, we scrambled either V0c loops 1.2 (L_1.2s_) or 3.4 (L_3.4s_) in the full-length protein and monitored interactions using an SPR-based method. V0c binding to immobilized complexin dissociates very slowly, indicating a strong interaction (Fig.1b). Binding of V0c containing a scrambled loop 3.4 to immobilized complexin was nearly identical to that of wild type V0c (Fig. 1b). However, V0c with scrambled loop 1.2 failed to bind complexin (Fig. 1b). The implication of V0c loop1.2 in complexin binding was confirmed by the experiment showing that a peptide corresponding to loop 1.2 totally inhibited V0c binding to immobilized complexin, while loop 3.4 peptide had no effect (Fig. 1c).

**Fig. 1.**
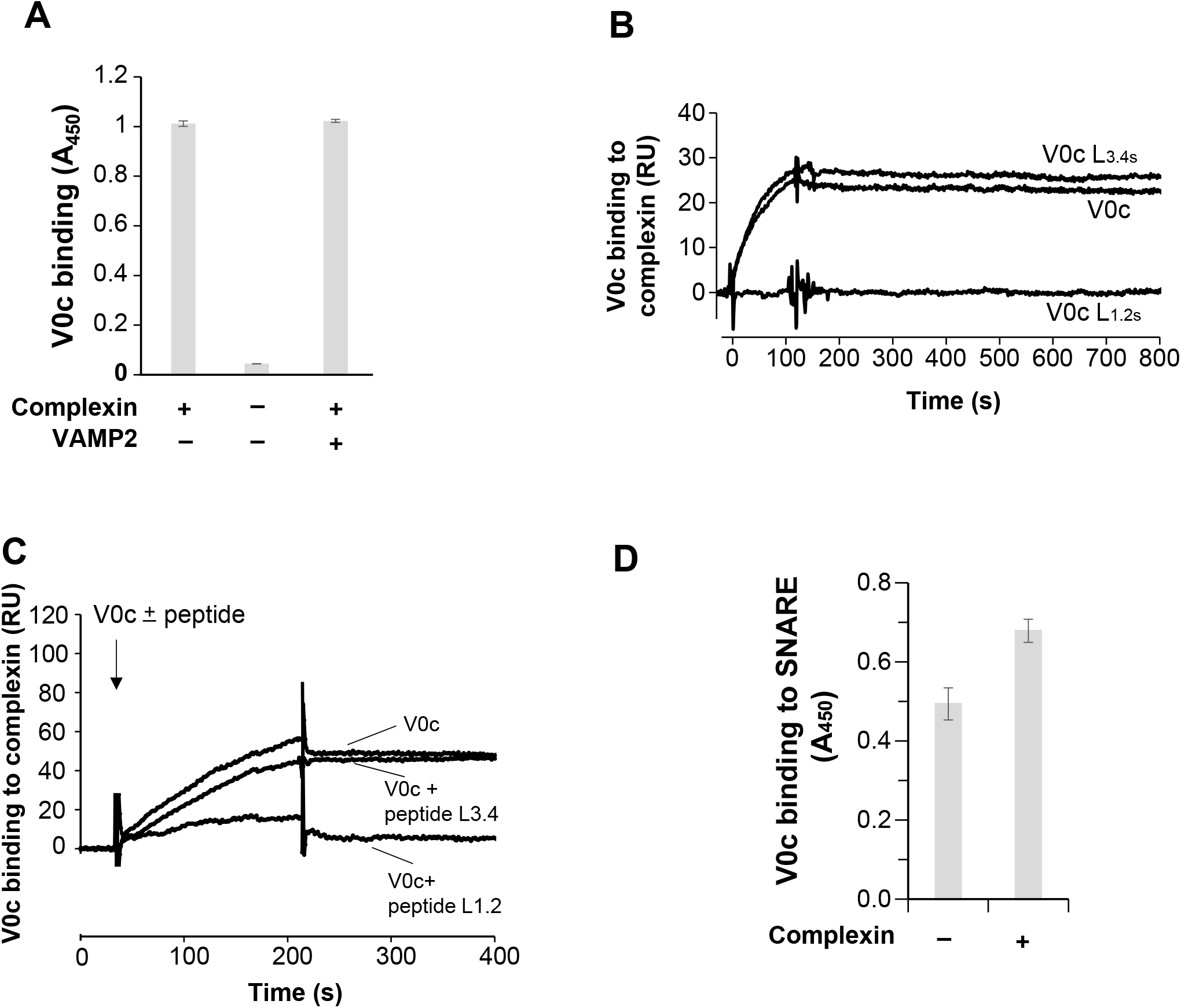
Characterisation of the V-ATPase V0c binding to complexin. **A** ELISA analysis of V0c binding to complexin. His-tagged complexin was immobilized or not in ELISA plates and binding of V0c (20 nM) was monitored by anti-HSV in the presence or absence of VAMP2 (100 nM). Note that V0c does not bind non-specifically BSA-saturated ELISA wells even at high concentrations (Supplemental. Fig. 2). Results are means of triplicate determination ± SD. Representative of 2 independent experiments **B** SPR measurement of V0c binding to complexin. V0c, V0c with scrambled loop 1.2 (V0c L1.2s) or scrambled loop 3.4 (V0c L3.4s) (50 nM) were injected for 2 min on a sensorchip functionalized with GST and GST-complexin. Background binding to GST has been subtracted from the presented results. Representative of 4 independent experiments. **C** V0c loop 1.2 peptide inhibits V0c binding to complexin. V0c (100 nM) was preincubated or not with L1.2 or L3.4 V0c peptides (10 µM) and injected on immobilized GST-complexin. Representative of 2 independent experiments **D** V0c binding to preassembled SNAREs. Purified preassembled (Supplemental Fig. 3) SNARE complex was immobilized in ELISA plates and binding of V0c (500 nM) was monitored by anti-HSV in the presence or absence of complexin (500 nM). Results are means of triplicate determination ± SD. Representative of 3 independent experiments

In accordance with the potentially late implication of V0c in exocytosis [33] [50], we investigated whether V0c interacts with the SNARE complex in the presence of complexin. The three SNARE proteins (devoid of any TM domains) were co-expressed in bacteria and SDS-resistant purified SNARE complexes were obtained (Supplemental Fig. 3). SNARE complexes were immobilized on ELISA plates and binding of V0c was assayed with or without preincubation with complexin. As shown in Fig.1d, V0c interacts with the SNARE complex in the presence of complexin. In summary, V0c interacts with complexin through the cytosolic loop 1.2 and the presence of complexin does not inhibit V0c binding to VAMP2 or to the SNARE complex.

### V0d interaction with V0c loop 1.2 precludes V0c association with complexin and the SNARE complex

Although V0d1 is a soluble protein, it remains stably associated with the V0 sector after dissociation of the V1 sector [51] [14] [15]. In order to assess the implication of V0d1 in the interaction of V0c with other molecular partners, we started by characterizing recombinant V0d1 binding to a series of immobilized proteins. As shown in Fig. 2a, ELISA experiments revealed that GST-V0d1 bound to immobilized V0c, but did not interact with VAMP2, the dimeric syntaxin/SNAP-25 t-SNARE complex, the trimeric SNARE complex, complexin or complexin associated with the SNARE complex.

**Fig. 2.**
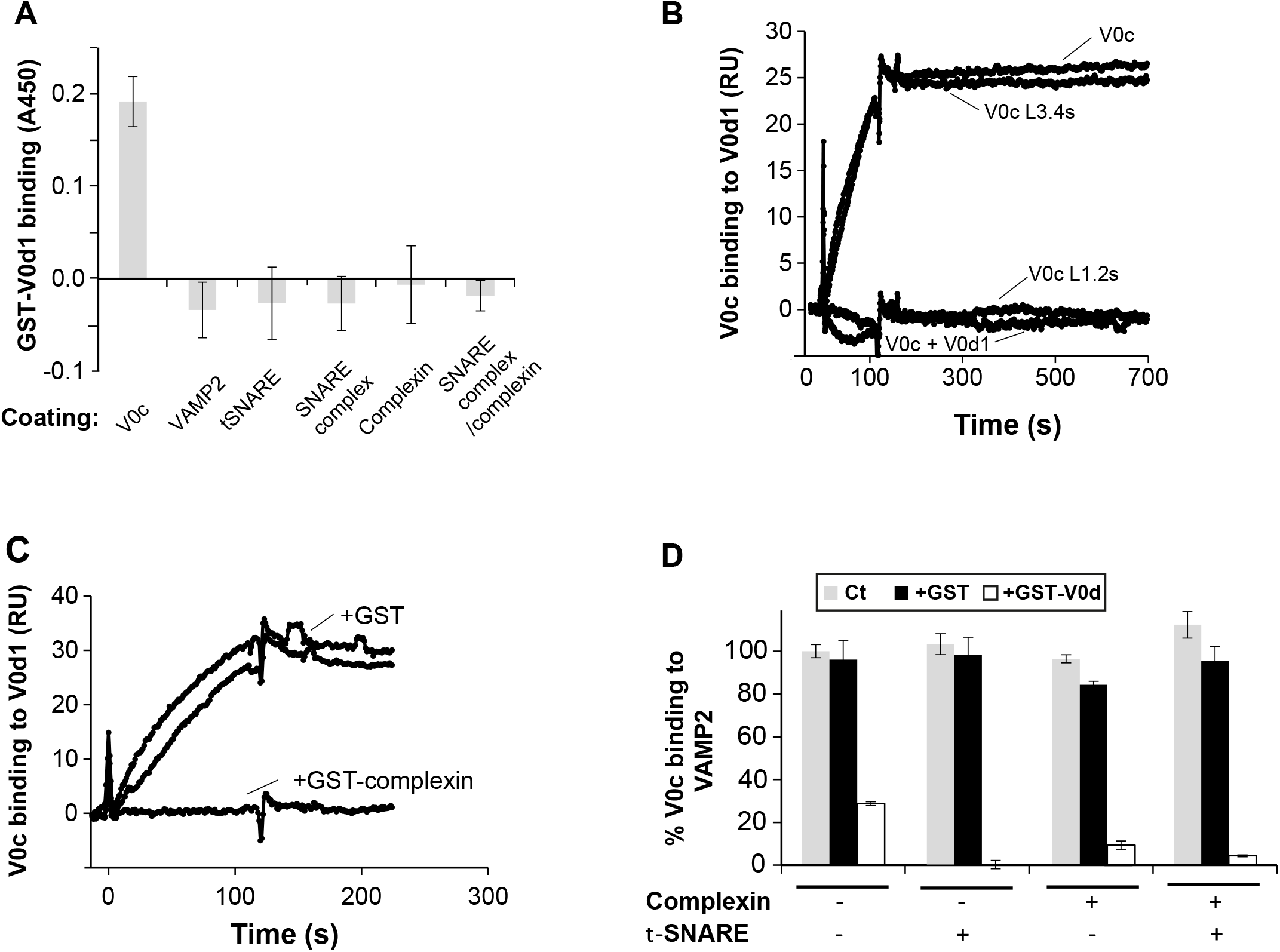
Characterisation of V0c/V0d1 interaction. **A** V0c, VAMP2, t-SNARE, complexin, the trimeric SNARE complex in the presence or absence of complexin were immobilized in ELISA plates and binding of GST-V0d1 and GST (50 nM) was measured. GST binding has been subtracted from the presented results. Representative of 3 independent experiments. **B** Recombinant wild type V0c (50 nM) ± 2 µM of soluble GST-V0d1, mutant V0c (V0c L1.2s or V0c L3.4s) containing respectively scrambled V0c loops 1.2 and 3.4 sequences were injected over consecutive flow cells of a SPR sensor-chip pre-functionalized respectively with GST and GST-V0d. Signals from GST coated flow cells have been subtracted **C** 50 nM recombinant wild type V0c was injected over consecutive flow cells of a SPR sensor-chip pre-functionalized with GST or GST-V0d1 in the presence of an excess of either GST or GST-complexin. Binding signals on precoated GST flow cells have been subtracted from represented sensorgrams. **D** Recombinant His-VAMP2 was immobilized in ELISA plates and V0c (20 nM) binding was monitored alone or in the presence of t-SNARE dimer (1 µM), complexin (0.2 µM) or a combination of both. For each combination, competition with 1 µM of either GST or GST-V0d1 were performed. Results are means of triplicate determinations ± SD. Representative of 3 independent experiments.

We then used SPR to identify the domain of V0c that interacts with V0d. As shown in Fig. 2b, purified recombinant V0c bound stably to immobilized V0d1 and showed no apparent dissociation, in accordance with previous data showing that V0d stably co-partitions with V0 in detergent-resistant membranes [11]. The specificity of this interaction was corroborated by the absence of V0c binding when it had been pre-incubated with an excess of V0d1 (Fig. 2b). Binding of V0c containing scrambled loop 3.4 to immobilized V0d1 was nearly undistinguishable from wild type V0c, however, V0c containing scrambled loop 1.2 failed to bind V0d1 (Fig. 2b). The involvement of V0c loop 1.2 in V0d binding was supported by the observation that the addition of free L_1.2_ peptide totally inhibited V0c binding to immobilized V0d1, while free L_3.4_ peptide addition had no effect (Supplemental Fig. 4). The involvement of V0c loop 1.2 in interaction with both V0d1 and complexin was confirmed by the observation that co-injecting a molar excess of complexin inhibited V0c binding to immobilized V0d1 (Fig 2c).

We then assessed if V0d could modulate V0c interaction with VAMP2 and the SNARE/complexin complex, using ELISA competition experiments. As shown in Fig. 2d, the presence of GST-V0d1 prevents V0c from binding to VAMP2 (inhibited by 71.3 % ± 3.2) and to in-situ reconstituted SNARE complex (inhibited by 100 %). The blocking effect of V0d is also observed in the presence of complexin for V0c binding to VAMP2 (inhibited by 90.7 % ± 1) and to the SNARE complex (inhibited by 96.1 % ± 6.2). This interference probably results from V0d interaction with V0c loop 1.2 inducing a steric hindrance of VAMP2 access to loop 3.4.

### Presynaptic V0d1 injection inhibits neurotransmission in SCG neurons

The importance of V0 subunits in the proton pump activity of the V-ATPase, and therefore in cell viability, compromises interpretation of data from gene inactivation and mRNA silencing to address the direct implication of these subunits in exocytosis. A multimer of V0c subunits constitutes the V-ATPase rotor, associated with a single V0d subunit. Although it does not have a membrane anchor, V0d behaves like an extrinsic membrane protein and V0d1 is not detected in cytosolic fractions (Supplemental Fig. 5). Based on our biochemical results indicating that V0d inhibits V0c interactions with VAMP2, SNARE complex or complexin, we reasoned that introduction of soluble V0d into the cell would stably occupy free binding sites on the multimeric V0c rotor, impeding association with exocytotic proteins. Superior cervical ganglion neurons (SCG) in culture have been widely used to explore the dynamics of presynaptic protein-protein interactions and understand their implication in neurotransmitter release [52]. As VAMP2 binding to V0c is involved in modulation of neurotransmission [22] and in order to test the hypothesis that V0d1 may regulate V0c interactions with SNARE proteins, we injected purified recombinant His-tagged V0d1 protein (6.5 µM) into presynaptic SCG neurons and monitored neurotransmitter release through postsynaptic recordings. As shown in Fig. 3a left, neurotransmission started to decrease a very few minutes after injection and reached a plateau shortly after 20 min with a mean inhibition ratio of 28% ± 3.9 p<0.01 at 9 min after injection and a maximal inhibition of 35% after 35 min. Of note control BSA injections did not lead to any significant decrease in neurotransmission (Fig. 3a right). As any perturbation of the V-ATPase function could drastically impact enzymatic activity and neurotransmitter uptake into synaptic vesicles, we verified whether the presence of excess V0d could impact proton pumping and consequently, if neurotransmission decrease does not result from a defect in acidification of the vesicle lumen, which may lead to reduced neurotransmitter loading. We therefore monitored synaptic vesicle V-ATPase activity in purified synaptic vesicles in the presence or absence of an excess of V0d1, by monitoring the intra-vesicular accumulation of fluorescent protonated Acridine Orange. As shown in Figure 3b, addition of up to 9 µM of V0d did not inhibit vesicular acidification in contrast to application of the proton pump inhibitor bafilomycin A1 which resulted in total inhibition of proton pump activity, and completely prevented Acridine Orange (AO) accumulation in synaptic vesicles (Fig. 3b). This clearly demonstrates that excess V0d does not perturb vesicular acidification. Hence a presynaptic V0d overload, which prevents V0c from binding to complexin and the SNARE complex rapidly diminished acetylcholine release by a mechanism independent of the proton pump activity of the V-ATPase.

**Fig. 3.**
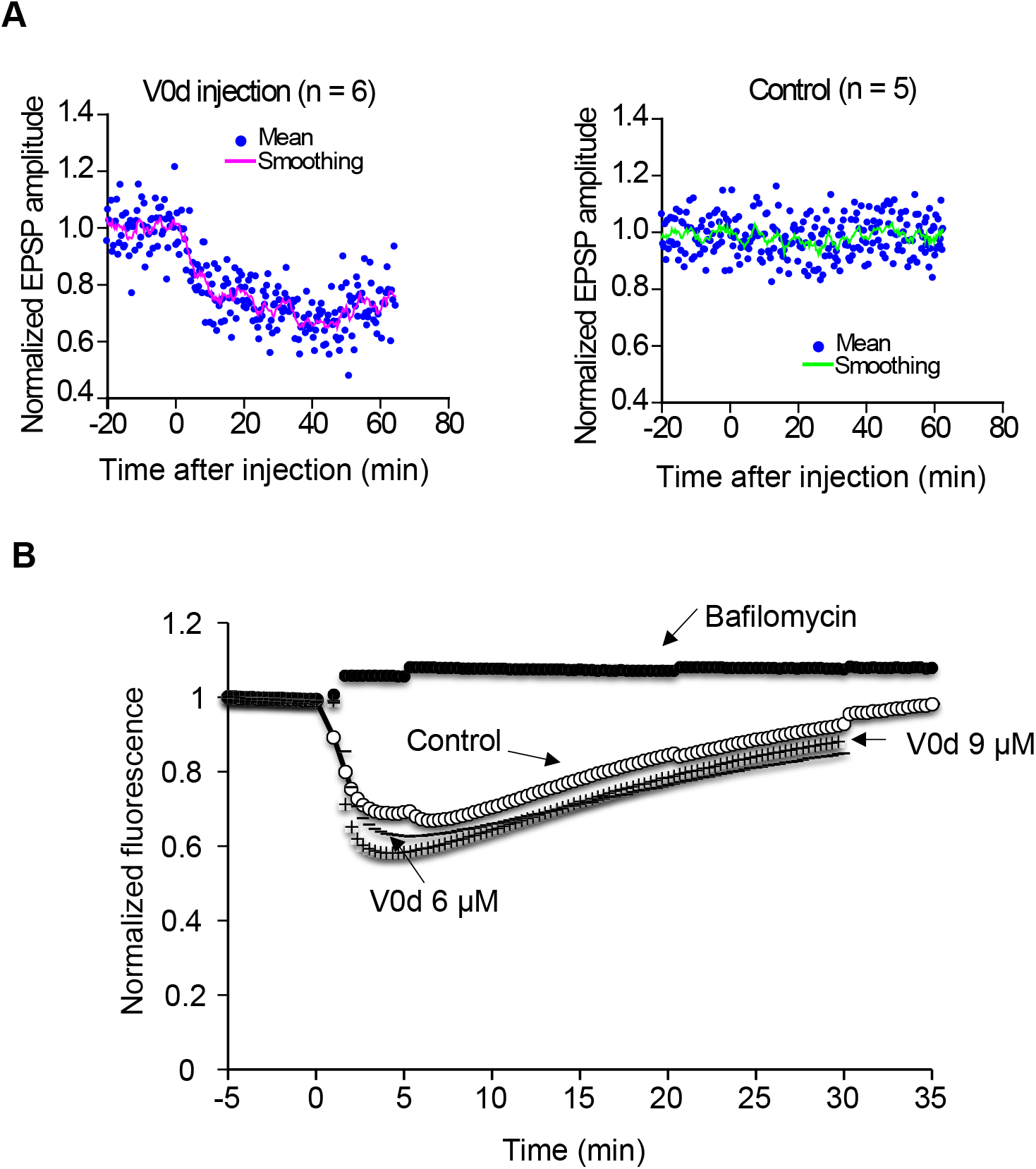
Effect of V0d injection on neurotransmission in SCG neurons. **A** Normalized excitatory postsynaptic potentials (EPSP) from connected superior cervical ganglion (SCG) neurons were plotted against time. EPSP were monitored at least 20 min before starting injections (Time = 0 min). Individual recording values are represented by dots. Smoothed values of normalized and averaged EPSP (unbroken line) were recorded every 10 sec in the presence of 6.5 µM His-HA-V0d (Left; n=6) and BSA (Right; n=5). **B** Differential effects of V0d1 and bafilomycin on synaptic vesicle proton pump activity and synaptic vesicle acidification. Acridine orange fluorescence was monitored over time in the absence or presence of 2 µM bafilomycin or two different concentrations of V0d (6 and 9 µM). The zero (min) point indicates addition of Mg^2+^/ATP. Results are representative of 3 independent experiments.

### V0d overexpression perturbs catecholamine release after in chromaffin cells

To investigate the effects of V0d1 on catecholamine exocytosis, we overexpressed myc-tagged V0d1 from a bicistronic mRNA co-expressing EGFP in chromaffin cells. Using carbon-fiber amperometry [53] [54], we monitored catecholamine release triggered by KCl depolarization and analyzed several exocytosis parameters (Fig. 4). Catecholamine release from EGFP-expressing cells was not different in any of the measured parameters compared to un-transfected cells (data not shown). Among all the parameters analyzed, four were significantly decreased upon V0d overexpression: spike number (57 ± 4 % of control), the total amount of currents during the spikes I_max_ (# 74 ± 2 % of control), the quantal size Q (70 ± 4 % of control) as well as the pre-spike foot (PSF) charge Q (# 73 ± 6 % of control). The kinetic parameters of individual exocytosis events T Half and the time to peak were not significantly affected. The percentage of spikes with a detectable foot (about 25% in the two conditions) was not modified and no significant variation in the pre-spike or foot parameters were observed. Altogether these results showed that V0d overexpression dramatically reduced the number of exocytotic events and reduced the amount of catecholamine released in the remaining events.

**Fig. 4.**
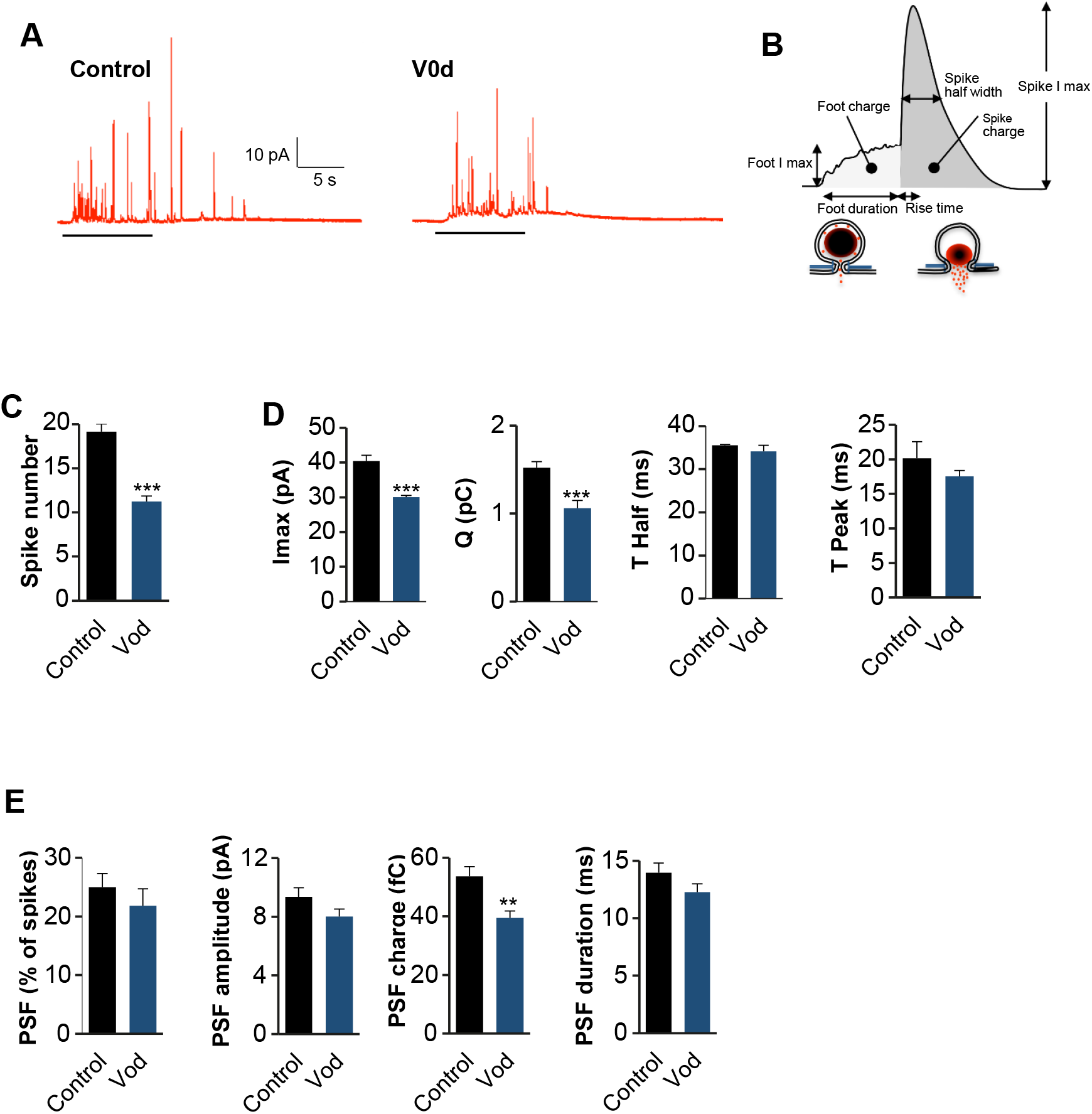
Effect of V0d1 overexpression on chromaffin cells exocytosis. **A** Bovine chromaffin cell in culture overexpressing EGFP alone (Control) or V0d1 / EGFP (V0d) were stimulated with a local application of 100 mM of K^+^ depolarizing solution for 10 s (bars bellow the curves) and catecholamine secretion was monitored using carbon fiber amperometry. Typical amperometric recordings are presented. **B** Scheme illustrating the different parameters of the amperometric spike. **C** The histogram corresponds to the total number of amperometric spikes recorded per cell in response to 10 s stimulation. **D** Amperometric spike parameters analyzed in Control or V0d expressing bovine chromaffin cells. Data are expressed as means ± S.D. (n > 60 cells for each condition from four independent cell cultures; * *p*<0.05, ** *p*<0.01, *** *p*<0.001 compared to control).

### V0c silencing reduces catecholamine release in chromaffin cells

Our biochemical analysis suggests that the effect of V0d1 is exerted through a reduction in the number of V0c interaction sites available for complexin/SNARE complexes. In order to corroborate this interpretation, we monitored catecholamine release from chromaffin cells and analyzed the parameters of exocytosis after siRNA silencing of V0c (Fig.5a, b). V0c silencing leading to a reduction of V0c levels by 40-50% without affecting V0a1 and V1A expression levels (data not shown) resulted in an approximately 40% decrease in the quantal content (Fig.5) presumably due to a perturbation of catecholamine loading into secretory granules. Interestingly, very similar to the effects of V0d1 overexpression, V0c silencing, using two distinct siRNA, significantly decreased spike number / cell (# 50% of control) and I_max_ (# 70% of control) (Fig. 5). These results suggest that V0c availability is implicated in determining the number of fusion events. In addition, and in contrast to V0d1 overexpression, V0c silencing led to a significant decrease in several additional PSF parameters: the number of PSF per cell (# 40% of control), the amount of current per PSF (# 80% of control), the quantal size Q of the foot (# 60% of control). In order to ascertain the specificity of the observed V0c siRNAs effects, we performed rescue experiments and we reintroduced wild type V0c insensitive to the siRNA. As shown in Fig. 5, rescue experiments restore wild type recording parameters demonstrating that the observed changes using the siRNA are due to a specific decrease of V0c.

To assess more specifically the effect of V0d1 overexpression and V0c silencing on catecholamine content in secretory granules we used the selective noradrenalin and dopamine fluorescent indicator NS510, which specifically accumulates in chromaffin secretory granules [55]. As illustrated in Supplemental Fig. 6 overexpression of V0d1 and silencing of V0c reduced by 16% and 30%, respectively, the mean intensity of the NS510 staining. These observations suggest that long term overexpression of V0d1, as well as a reduction in the level of V0c expression, moderately reduced catecholamine content in secretory granules that might result from a reduction of secretory granule acidification. It is however unlikely that this moderate effect on catecholamine loading could explain the potent reduction in the number of individual exocytotic events, arguing for the importance of the V0d/V0c interaction in the modulation of exocytosis.

**Fig. 5.**
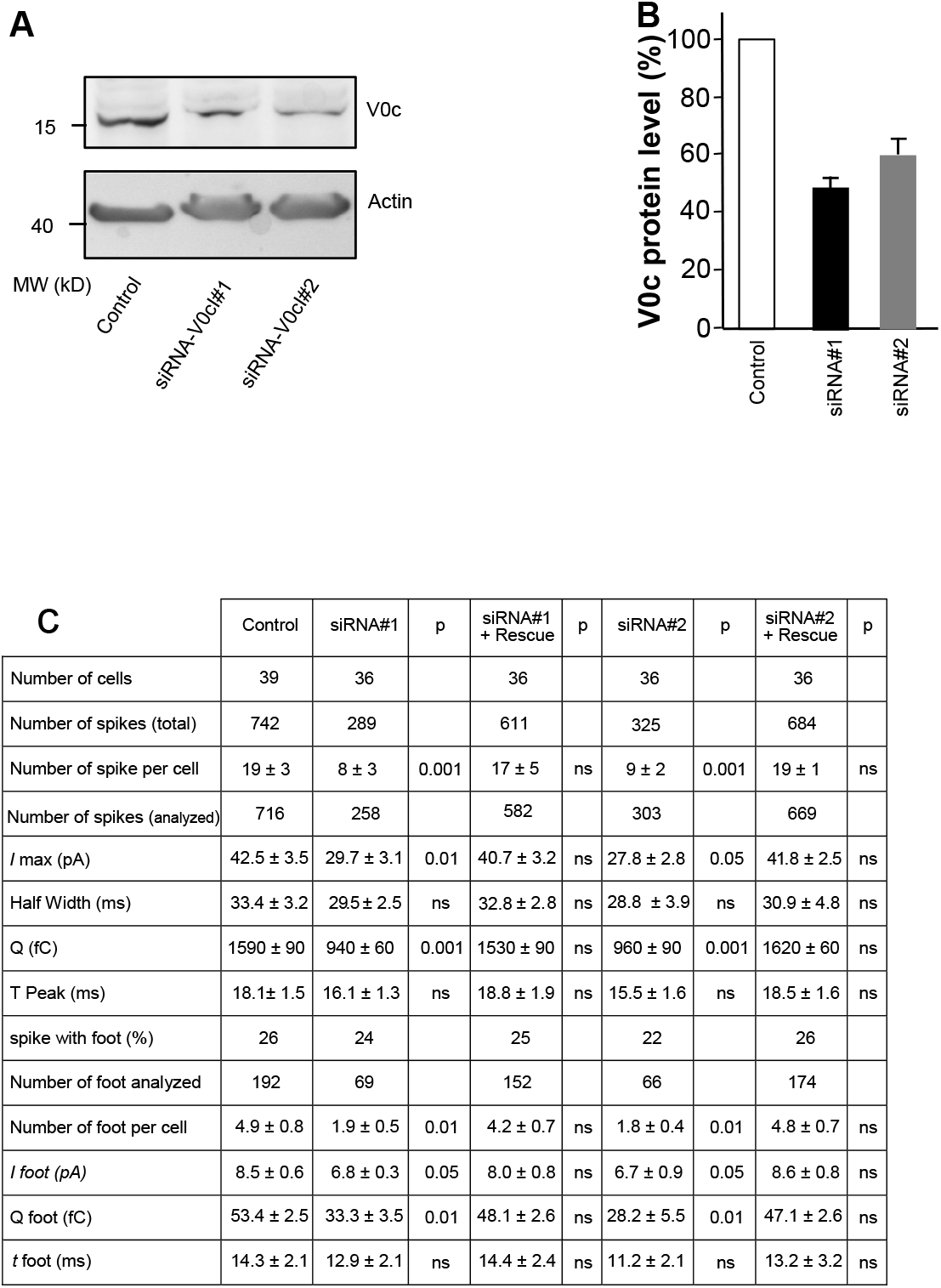
Effect of V0c silencing on catecholamine release from chromaffin cells. **A** Two different siRNAs targeting bovine V0c (Thermo Fisher Scientific) reduced V0c expression level after 96 h. Actin served as a loading control. **B** Quantification of the silencing effect on V0c expression (n = 6). Data are means ± SEM. **C** Bovine chromaffin cells nucleoporated with the indicated V0c siRNA and coexpressing GFP or rat V0c (siRNA resistant) and GFP. Bars, 5 µm. 96 h after nucleofection, the two siRNAs targeting V0c significantly affected various amperometric signal parameters when catecholamine release was evoked by a 10 s application of 100 mM KCl. Co-expression of a siRNA-resistant V0c rescued all parameters to the control condition.

## Discussion

It has been reported that the V1 domain of the V-ATPase is absent from fully loaded as well as docked synaptic vesicles [7] [6] and that conformational rearrangements in V0a and V0d subunits take place upon V1 dissociation leading to a so-called auto-inhibited V0 domain [14] [15] [16] [5]. Several reports already showed that independently of its participation in the proton pumping activity of the V-ATPase, the V0 domain is implicated in exocytosis [21] [22] [23] as well as in membrane fusion of intracellular compartments [24] [25] [26] [27]. In addition, direct interactions of V0 domain subunits with the minimal membrane fusion machinery, i. e. the SNARE complex or individual SNARE proteins, have been reported with functional consequences [56] [33] [31] [22]. Complexin is a major regulatory partner of the SNARE complex [49]. In vertebrates, among the 4 isoforms, complexin 1 and 2 are mainly neuronal and complexin 3, 4 are retina specific [57]. Despite its small size, complexin has a very intricate function with distinct adjacent and functionally different domains. Each of these domains can separately exert opposing properties as both an inhibitor or a facilitator of synaptic vesicle fusion modulating evoked and spontaneous release [58] [29] [49] [59] [60]. It interacts through its central α-helix domain with the SNARE complex [61] [62] [29] and is implicated in organizing high order fusogenic SNARE/synaptotagmin complexes [29] [63]. Structural analysis has shown that complexin is associated with the SNARE complex in an anti-parallel orientation, leaving the C-terminal domain oriented towards the N-terminal domains of syntaxin and VAMP in the assembled SNARE complex [61] [64]. Also, the interaction of the C-terminal domains of complexins 1 and 2 with lipids [65] and with the highly curved membranes of synaptic vesicles was suggested to be important for its inhibitory function [66] [67] [68].

In this study we report a direct interaction of the V0c subunit of the V-ATPase with complexin as well as the tetrameric complexin / SNARE complex. We show that the interaction of complexin with V0c is mediated by the loop 1.2. As the loop 3.4 of V0c binds VAMP2 and V0c binding to complexin is not competed by VAMP2, one may assume that V0c can interact concomitantly with two essential components of the exocytotic machinery. Moreover, we demonstrate that the loop 1.2 of V0c mediates an extremely stable interaction with V0d1 in line with the existence of contact sites between V0d and V0c rotor in yeast [15] and mammalian [18] V-ATPase. Our biochemical experiments indicate that an excess of V0d1 hinders V0c interaction with VAMP2, complexin and the complexin/SNARE complex. Since V0c interacts with complexin and VAMP2 via two distinct loops (loop 1.2 and loop 3.4 respectively) and complexin does not inhibit V0c/VAMP2 interaction, the inhibitory effect of V0d on V0c binding to VAMP2 may result from a steric hindrance due to the higher molecular weight of V0d compared to complexin. In order to get an insight into the physiological implication of V0c loop 1.2 in exocytosis, we reasoned that introducing soluble V0d would impede V0c loop 1.2 interaction with complexin and SNARE complex. In a first approach we monitored excitatory postsynaptic potentials (EPSPs) in connected superior cervical ganglion (SCG) neurons after presynaptic injection of bacterially-expressed and purified V0d1. SCG neurons are a well-established culture system suited for the study of neurotransmitter release mechanisms. The very short axonal connections provide a setup where somatically-injected effectors rapidly reach nerve terminals. Upon V0d1 microinjection in SCG neurons, we observed rapid inhibition of neurotransmission similar to that observed upon injection of a peptide corresponding to V0c L3.4 loop [33] and other agents that affect late steps in exocytosis [69]. As V0d is a functional component of the V-ATPase complex, we addressed the possibility that introducing free V0d could modify intramolecular interactions between V-ATPase subunits and consequently inhibit proton pumping, which would then compromise neurotransmitter loading into synaptic vesicles. In this case vesicles devoid of neurotransmitter might still fuse without generating an EPSP. To explore this possibility, we studied the effect of an excess of V0d1 on synaptic vesicle acidification using an Acridine Orange-based assay. Although the V-ATPase inhibitor bafilomycin produced an immediate and complete block of proton accumulation in synaptic vesicles, V0d1 concentrations higher than those injected into SCG neurons failed to decrease acidification. These data strongly suggest that an excess of cytosolic V0d1 is unlikely to perturb synaptic vesicle loading with neurotransmitters. Similar to inhibition of acetylcholine release in SCG neurons, catecholamine secretion was altered after V0d1 overexpression in chromaffin cells, a well-defined cellular model to study neuroendocrine secretion [70]. In addition to a significant decrease in the number of spikes (Fig. 4a, c) and maximal current reduction (Fig. 4d), the quantal content of granules was affected upon V0d1 overexpression (Fig. 4d) without affecting time to peak kinetics. Apart from the pre-spike foot (PSF) charge which was decreased, V0d1 overexpression did not affect any other PSF parameter (Fig. 4e). It is, however, of note that under these conditions of V0d1 overexpression a reduction of catecholamine content of approximatively 16% was observed using the fluorescent indicator NS510 (Supplemental Fig.6). The observed decrease in the quantal content by roughly 30% may thus partly be due to the long-term genetic nature of V0d1 overexpression. It is possible that co-translational abundance of V0d1 with endogenous native levels of other V-ATPase subunits may perturb V-ATPase assembly and therefore granule acidification and catecholamine loading. On the other hand, the modest effect observed on catecholamine loading under these conditions is unlikely to explain the robust differences in spike number, Imax and charge observed when V0d1 is overexpressed, arguing for a role of the V0d1-V0c interaction in the regulation of exocytosis.

In an earlier report, we have shown that inhibiting V0c interaction with VAMP2 significantly inhibits neurotransmission [33]. Previous data showed that the native multimeric V0c rotor binds to a single V0d1 subunit [1] [5] [18]. Our electrophysiological recordings in SCG neurons as well as the biochemical results suggest that inhibition of exocytosis upon V0d1 injection is likely due to binding of the exogenous V0d1 to V0c subunits in the V0 rotor that are not in contact with the single intrinsic V0d1 subunit. This consequently could prevent V0c interaction with complexin and SNARE proteins. In order to corroborate this hypothesis in chromaffin cells and compare the effect of V0d1 overexpression and V0c availability on catecholamine secretion, we performed siRNA V0c downregulation (Fig. 5 a, b) and monitored catecholamine secretion parameters. As expected, the granule quantal content was significantly decreased but catecholamine secretion, although diminished, still took place (Fig. 5c). Both V0c siRNAs triggered a decrease in V0c expression and lead to an inhibition of the number of individual release events as well as maximal current that reached very similar levels of inhibition that were observed upon V0d1 overexpression. In contrast to the effects of V0d1 overexpression, a reduced V0c expression level affected the number of PSF per cell, the spike shape (*I*max and half width), the foot*I*max highlighting the importance of V0c availability in modulating release. Previous structural studies of the yeast V-ATPase showed that, in the auto-inhibited V0 domain, V0d interacts with the N-terminal domain of V0a upon V1 dissociation and engages interactions with the V0c rotamer subunits [14] [15] [16] [5]. In bovine V-ATPase, V0d show stable interactions with 4 c-subunits and 1 c” in the assemble V-ATPase [18]. However, how the exogenous V0d incorporates into auto-inhibited V0 is still to be determined at the structural level.

All these data reinforce our understanding of the implication of the V-ATPase V0 subunits in modulating SNARE-dependent neurotransmission and corroborate the importance of V0c in SNARE-dependent neurosecretion. In a fully assembled V-ATPase, a V0c multimer accommodates only one V0d subunit [1] [5]. Moreover, we show that the V0d subunit is absent from cytosol. We speculate that, upon V1 dissociation, V0c subunits that are not linked to V0d1 would be free to bind VAMP2 as well as the cytosolic complexin [71]. The V0c subunits of the V0 sector associated with several VAMP2 molecules would interact with membrane t-SNAREs and engage the formation of a SNARE rosette around the V0c rotor [72]. Upon overexpression of V0d1, V0d1-free V0c subunits in a V0 rotor may become sequestered and hindered from binding to VAMP2 [33] and complexin resulting in an inhibition of formation of the SNARE rosette and thereby neurotransmitter release. Using molecular dynamics [73], it has recently been suggested, that confinement of a rosette of SNARE complexes [72] [74] is essential for rapid fusion pore formation, and fusion pore expansion is accompanied by a release from this confinement. A potential consequence of the presence of an excess of V0d1 would be that V0c would fail to form the SNARE complex rosette (Fig. 6) leading to a decrease in the frequency of membrane fusion events. Whether the single endogenous V0d1 subunit per V-ATPase is implicated in modulating exocytosis has not yet been addressed, future investigations are needed to clarify this point.

**Fig. 6.**
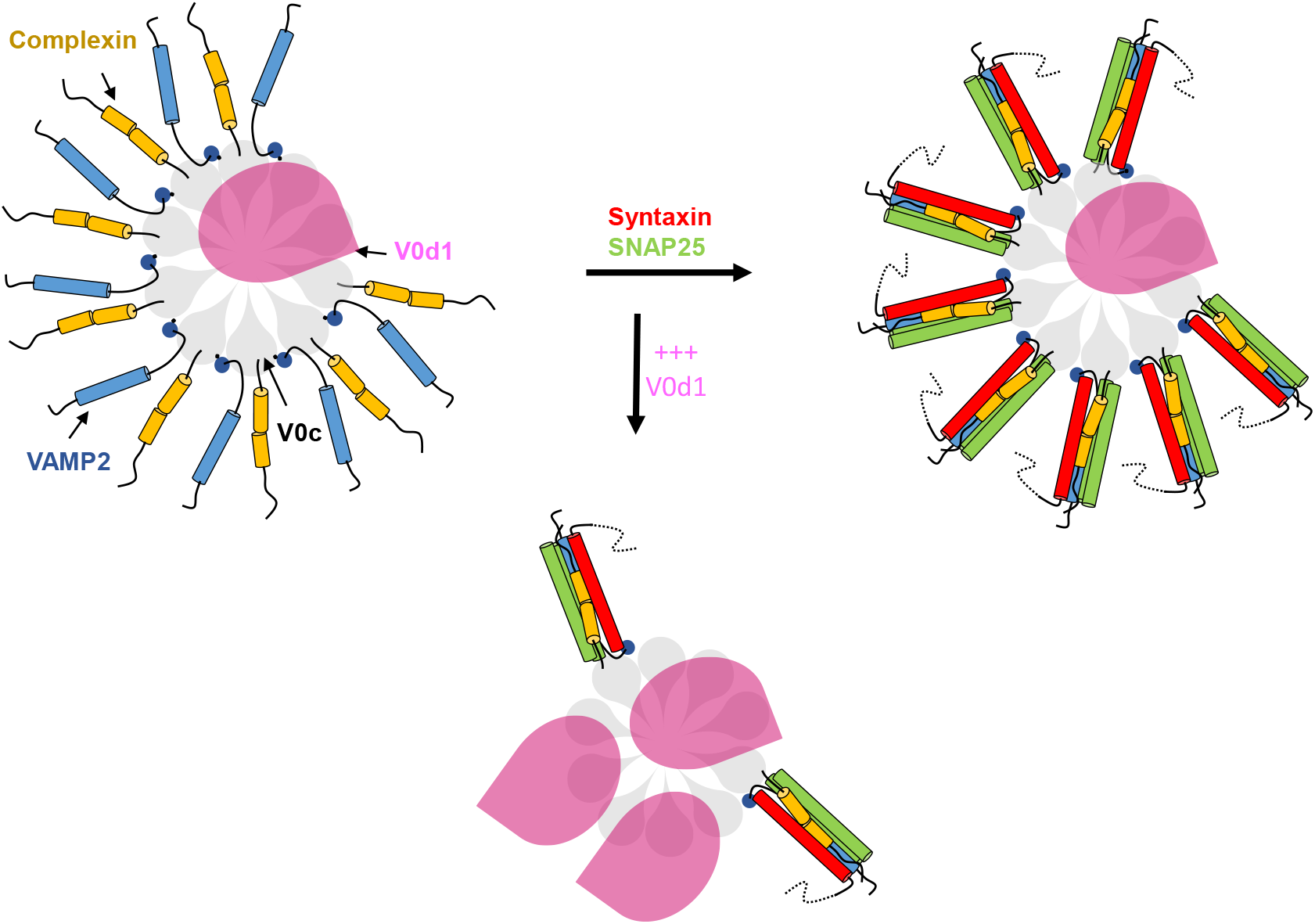
Speculative model of the SNARE-rosette organising role of the V-ATPase V0c rotor. **A** Auto-inhibited V0 rotor in the membrane of a fully loaded synaptic vesicle may locally concentrate VAMP2 and complexin. **B** Upon synaptic vesicle docking, the rotor of V0c / VAMP2 / complexin may serve as an organizer around which a rosette of SNARE would assemble. **C** In the presence of an excess of V0d, the capacity of V0c to organize the VAMP2 / complexin rosette and therefore SNARE complexes would be diminished thus affecting efficient exocytosis.

## Abbreviations

CALI: Chromophore-assisted light inactivation FC12 Fos-Choline-12
NS510: Neurosensor 510
SCG: Superior cervical ganglion
SDS: Sodium dodecyl sulfate
SPR: Surface plasmon resonance
TMD: Transmembrane domain

## Author contributions

OEF, NV, CL, and MSeagar conceived the study. YM, QW, MR, YF, CI and MSangiardi performed biochemical experiments. CL performed SPR experiments. YM, FY and MSangiardi performed expression plasmids preparation and preliminary expression tests. SM performed SCG recordings. QW and MR performed experiments on chromaffin cells. OEF, CL, NV and MSeagar wrote the manuscript. All authors edited and reviewed the manuscript.

## Acknowledgments

We thank Lea RODRIGUEZ for preliminary biochemical experiments

## Funding

This work was supported by INSERM, the CNRS and the ANR (Agence Nationale pour la Recherche grant ANR-11-BSV2-020-MOMENT Blanc SVSE2 to O.E.F. and grant ANR-19-CE44-0019 to N.V.)

## Data availability

Not applicable

## Declarations Conflict of interest

The authors declare that they have no conflict of interest

## Ethical approval

Not applicable

## Consent to participate

All authors approved submission

## Consent to publish

All authors approved publication

## Electronic supplementary material

**Supplemental Fig 1.**
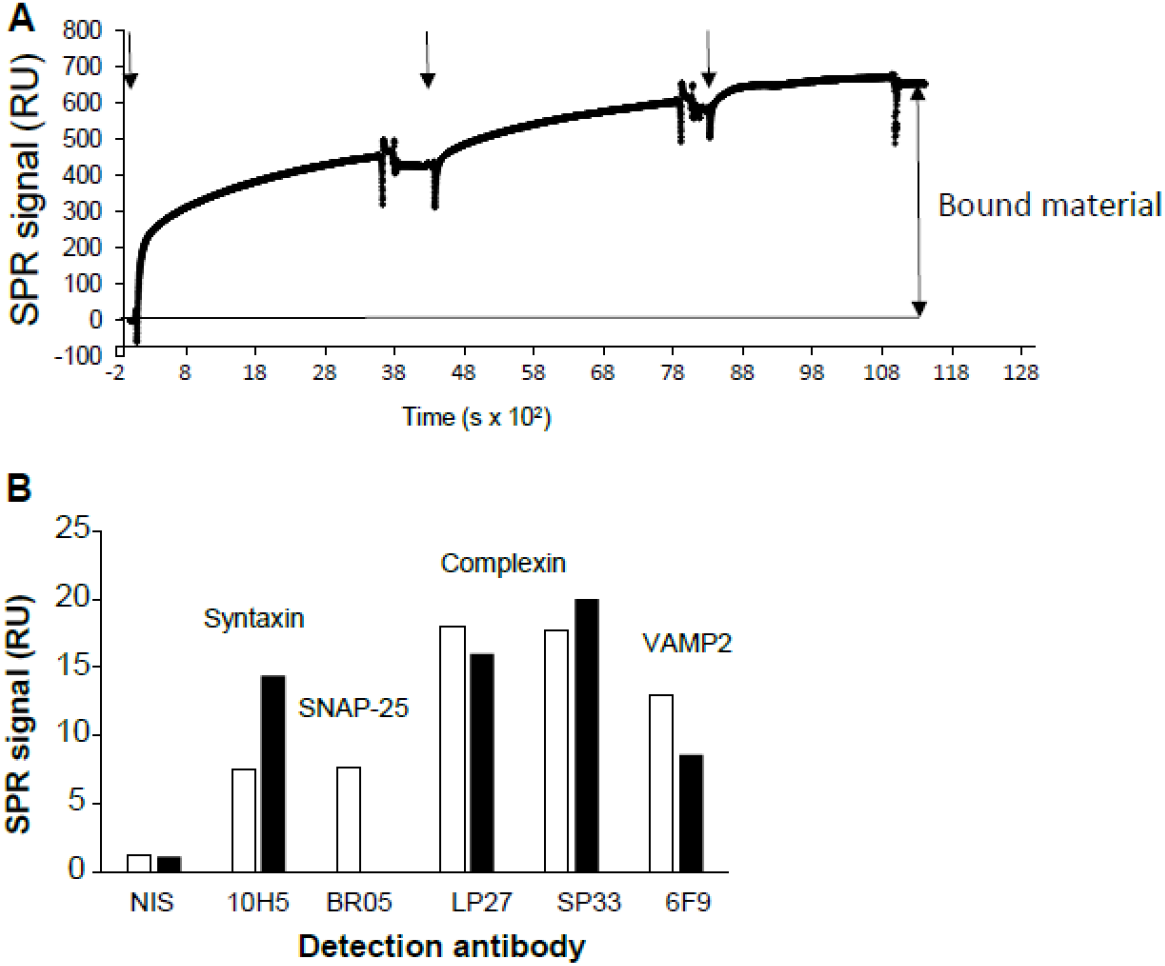
V0c captures SNARE proteins and complexin from rat brain extract. **A** Example of specific immobilization of solubilized rat brain proteins on V0c using SPR: recombinant HSV-tagged V0c was captured (1000 RU) on a sensor chip using anti-HSV antibodies. LP1 rat brain extract was injected 3 times (arrows) at 1 µl/min into the flow cell. This injection yields about 600 RU of stable and specific binding compared to a control flow cell loaded with only anti-HSV. Background in the absence of V0c has been subtracted from the presented data. **B** Specific signal resulting from injection of non-immune (NIS), anti-syntaxin1 (10H5), anti-SNAP25 (Br05), anti-complexin (LP27, SP13) and anti-VAMP2 (6F9) antibodies over brain extract proteins captured on immobilized V0c as illustrated in **A**. Detection levels (RU) are represented in histograms. Open and black bars are two independent experiments.

**Supplemental Fig 2.**
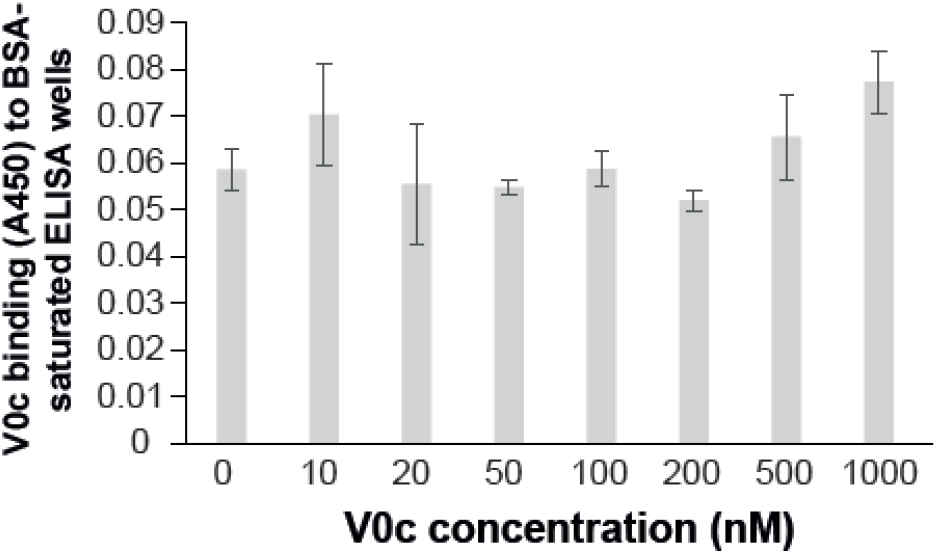
Background levels of V0c binding in ELISA plates. V0c binding to BSA-saturated ELISA wells was probed using increasing concentrations of V0c (10-1000 nM). Binding was detected using anti-HSV antibodies. Note that binding signals were equivalent to a condition in which V0c was not added. Results are means of triplicate determination ± SD.

**Supplemental Fig 3.**
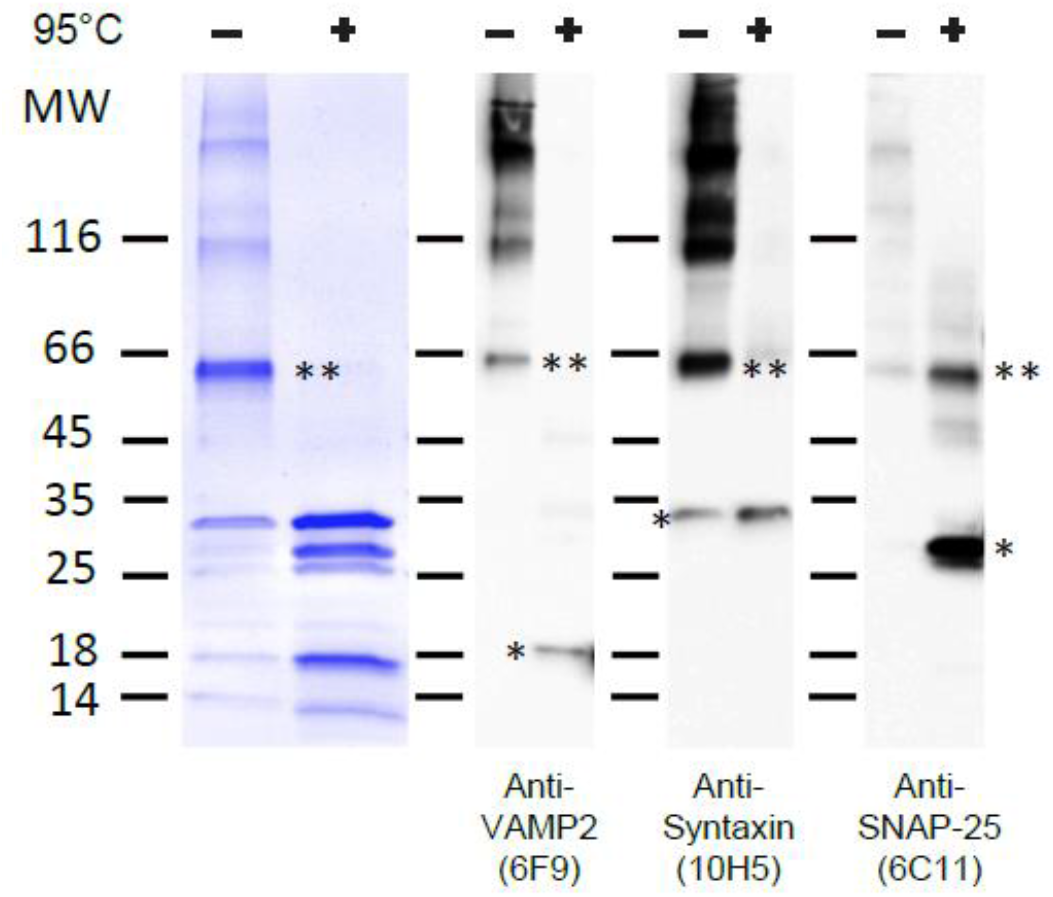
Quality control of SNARE complex formation. 4 µg of the purified SNARE complex were either heated or not at 95°C in sample buffer and analysed by SDS PAGE and Coomassie blue staining (left). The molecular identity of the components of this complex was verified by Western blot using anti-VAMP2 (6F9), anti-syntaxin1(10H5) and anti-SNAP-25 (6C11) antibodies (right). Note that the SNAP-25 epitope recognized by 6C11 is mostly hidden in the assembled SNARE complex. Positions of the SNARE monomers and the SNARE complex are indicated respectively by (*) and (**).

**Supplemental Fig 4.**
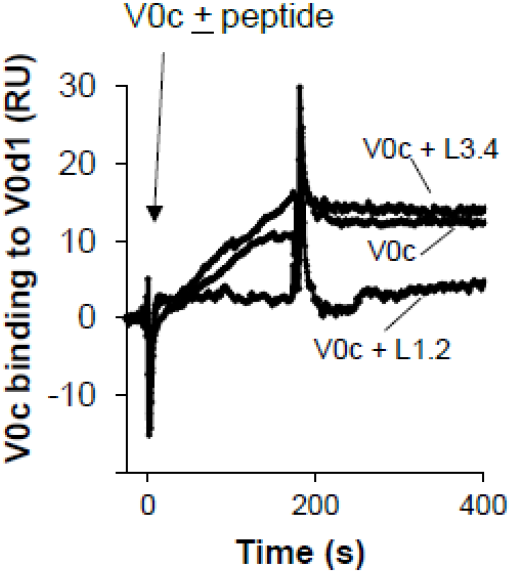
Characterisation of V0c/V0d1 interaction. 50 nM of recombinant wild type V0c was injected over consecutive flow cells of an SPR sensorchip pre-functionnalized with GST or GST-V0d1 in the presence of an excess of either V0c loop 1.2 (L1.2) or loop 3.4 peptides (L3.4). Binding signals on precoated GST flow cells have been subtracted from represented sensorgrams.

**Supplemental Fig 5.**
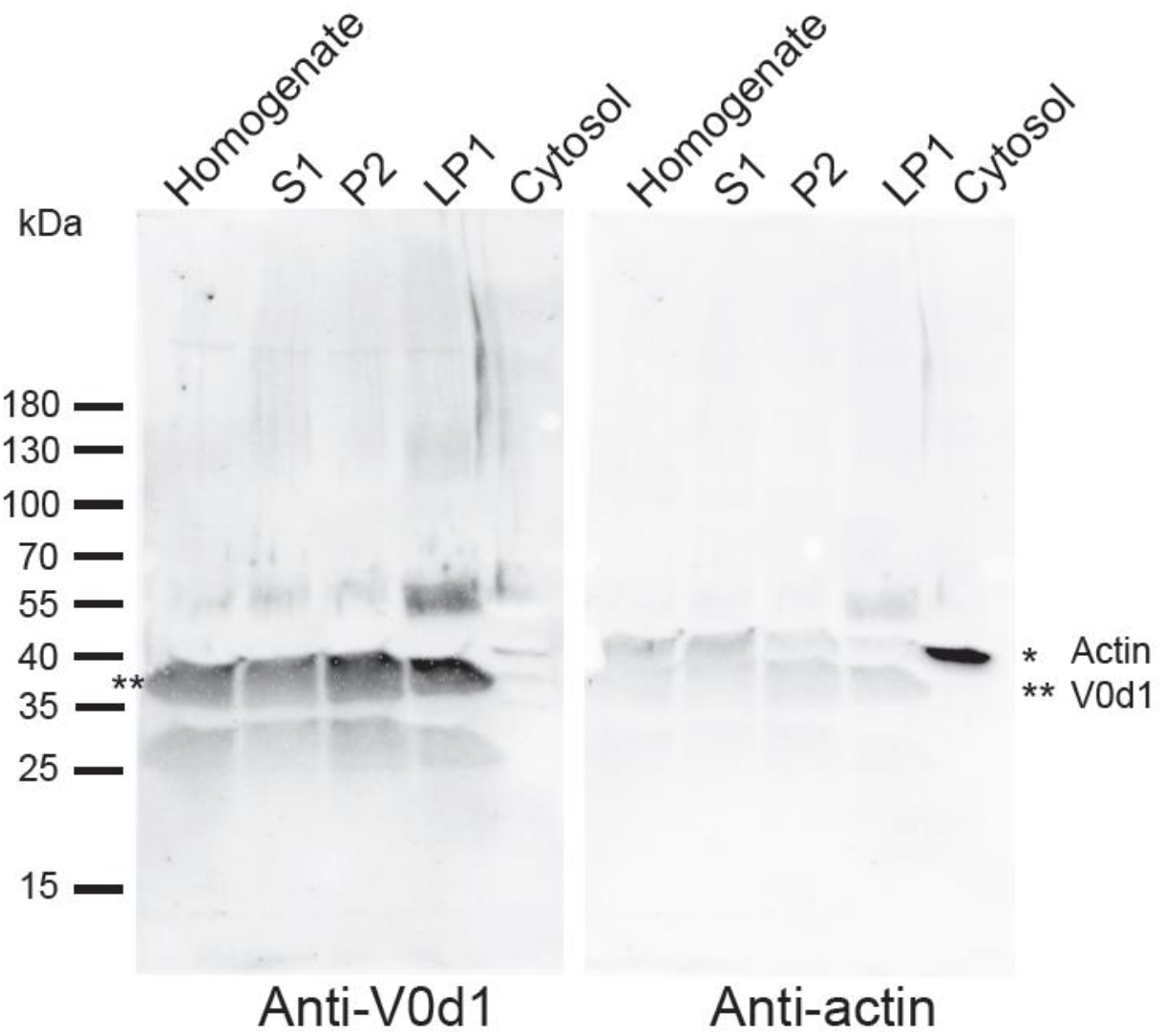
Distribution of V0d1 in different membrane fractions. The distribution of V0d1 was analysed by western blot in different membrane fractions of mice brain lysates (30 μg / well). (* = actin; ** = V0d1). Note the absence of V0d1 from the cytosolic fraction.

**Supplemental Fig 6.**
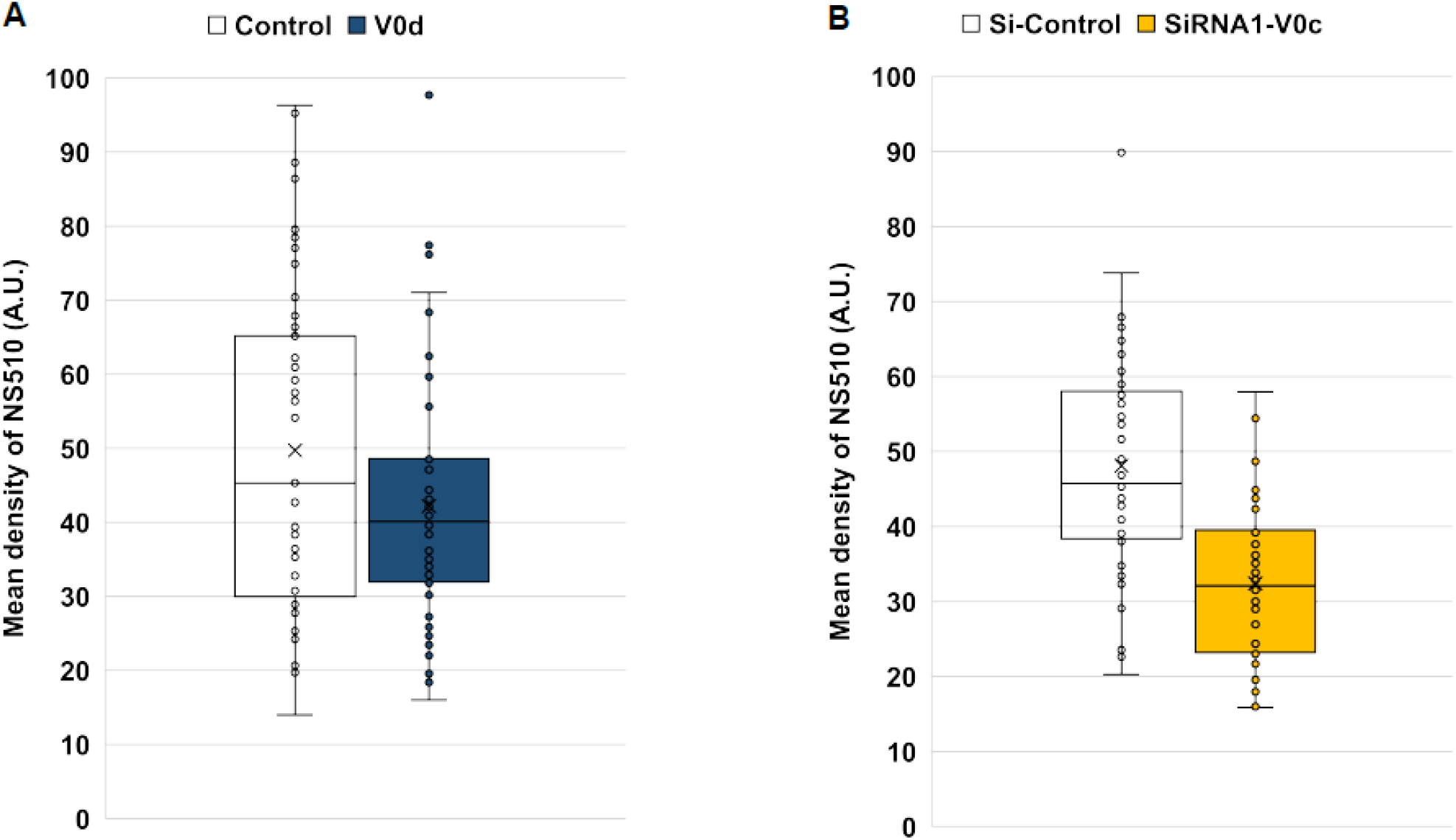
Effect of V0d overexpression and V0c silencing on catecholamine loading in secretory granules of bovine chromaffin cells. Chromaffin cells overexpressing V0d and TurboRFP (**A**) or silenced for V0c (**B)** were incubated with NS510. Quantification of accumulated NS510 signal was performed from individual cells (n=53) from three independent experiments for each condition.

## Notes

### Competing Interest Statement

The authors have declared no competing interest.

## References

1. Forgac M (2007) Vacuolar ATPases: rotary proton pumps in physiology and pathophysiology. Nat Rev Mol Cell Biol 8 (11):917–929. doi:10.1038/nrm2272

2. Hurtado-Lorenzo A, Skinner M, El Annan J, Futai M, Sun-Wada GH, Bourgoin S, Casanova J, Wildeman A, Bechoua S, Ausiello DA, Brown D, Marshansky V (2006) V-ATPase interacts with ARNO and Arf6 in early endosomes and regulates the protein degradative pathway. Nature cell biology 8 (2):124–136

3. Yokoyama K, Nakano M, Imamura H, Yoshida M, Tamakoshi M (2003) Rotation of the proteolipid ring in the V-ATPase. J Biol Chem 278 (27):24255–24258

4. Toei M, Saum R, Forgac M (2010) Regulation and Isoform Function of the V-ATPases. Biochemistry 49 (23):4715–4723

5. Abbas YM, Wu D, Bueler SA, Robinson CV, Rubinstein JL (2020) Structure of V-ATPase from the mammalian brain. Science 367 (6483):1240–1246. doi:10.1126/science.aaz2924

6. Bodzeta A, Kahms M, Klingauf J (2017) The Presynaptic v-ATPase Reversibly Disassembles and Thereby Modulates Exocytosis but Is Not Part of the Fusion Machinery. Cell Rep 20 (6):1348–1359. doi:10.1016/j.celrep.2017.07.040

7. Poea-Guyon S, Ammar MR, Erard M, Amar M, Moreau AW, Fossier P, Gleize V, Vitale N, Morel N (2013) The V-ATPase membrane domain is a sensor of granular pH that controls the exocytotic machinery. J Cell Biol 203 (2):283–298. doi:10.1083/jcb.201303104

8. Thaker YR, Roessle M, Gruber G (2007) The boxing glove shape of subunit d of the yeast V-ATPase in solution and the importance of disulfide formation for folding of this protein. J Bioenerg Biomembr 39 (4):275–289. doi:10.1007/s10863-007-9089-7

9. Smith AN, Francis RW, Sorrell SL, Karet FE (2008) The d subunit plays a central role in human vacuolar H(+)-ATPases. J Bioenerg Biomembr 40 (4):371–380. doi:10.1007/s10863-008-9161-y

10. Zhang J, Myers M, Forgac M (1992) Characterization of the V0 domain of the coated vesicle (H+)-ATPase. J Biol Chem 267 (14):9773–9778

11. Lafourcade C, Sobo K, Kieffer-Jaquinod S, Garin J, van der Goot FG (2008) Regulation of the V-ATPase along the endocytic pathway occurs through reversible subunit association and membrane localization. PLoS One 3 (7):e2758. doi:10.1371/journal.pone.0002758

12. Bauerle C, Ho MN, Lindorfer MA, Stevens TH (1993) The Saccharomyces cerevisiae VMA6 gene encodes the 36-kDa subunit of the vacuolar H(+)-ATPase membrane sector. J Biol Chem 268 (17):12749–12757

13. Thaker YR, Hunke C, Yau YH, Shochat SG, Li Y, GrÃ¼ber G (2009) Association of the eukaryotic V1VO ATPase subunits a with d and d with A. FEBS Letters 583 (7):1090–1095

14. Mazhab-Jafari MT, Rohou A, Schmidt C, Bueler SA, Benlekbir S, Robinson CV, Rubinstein JL (2016) Atomic model for the membrane-embedded VO motor of a eukaryotic V-ATPase. Nature 539 (7627):118–122. doi:10.1038/nature19828

15. Roh SH, Stam NJ, Hryc CF, Couoh-Cardel S, Pintilie G, Chiu W, Wilkens S (2018) The 3.5-A CryoEM Structure of Nanodisc-Reconstituted Yeast Vacuolar ATPase Vo Proton Channel. Mol Cell 69 (6):993–1004 e1003. doi:10.1016/j.molcel.2018.02.006

16. Vasanthakumar T, Bueler SA, Wu D, Beilsten-Edmands V, Robinson CV, Rubinstein JL (2019) Structural comparison of the vacuolar and Golgi V-ATPases from Saccharomyces cerevisiae. Proc Natl Acad Sci U S A 116 (15):7272–7277. doi:10.1073/pnas.1814818116

17. Smith AN, Borthwick KJ, Karet FE (2002) Molecular cloning and characterization of novel tissue-specific isoforms of the human vacuolar H(+)-ATPase C, G and d subunits, and their evaluation in autosomal recessive distal renal tubular acidosis. Gene 297 (1-2):169–177

18. Wang R, Long T, Hassan A, Wang J, Sun Y, Xie XS, Li X (2020) Cryo-EM structures of intact V-ATPase from bovine brain. Nat Commun 11 (1):3921. doi:10.1038/s41467-020-17762-9

19. Miura GI, Froelick GJ, Marsh DJ, Stark KL, Palmiter RD (2003) The d subunit of the vacuolar ATPase (Atp6d) is essential for embryonic development. Transgenic Res 12 (1):131–133

20. Nishi T, Kawasaki-Nishi S, Forgac M (2003) Expression and function of the mouse V-ATPase d subunit isoforms. J Biol Chem 278 (47):46396–46402. doi:10.1074/jbc.M303924200

21. Liegeois S, Benedetto A, Garnier JM, Schwab Y, Labouesse M (2006) The V0-ATPase mediates apical secretion of exosomes containing Hedgehog-related proteins in Caenorhabditis elegans. J Cell Biol 173 (6):949–961. doi:10.1083/jcb.200511072

22. Hiesinger PR, Fayyazuddin A, Mehta SQ, Rosenmund T, Schulze KL, Zhai RG, Verstreken P, Cao Y, Zhou Y, Kunz J, Bellen HJ (2005) The v-ATPase V0 subunit a1 is required for a late step in synaptic vesicle exocytosis in Drosophila. Cell 121 (4):607–620. doi:10.1016/j.cell.2005.03.012

23. Wang D, Epstein D, Khalaf O, Srinivasan S, Williamson WR, Fayyazuddin A, Quiocho FA, Hiesinger PR (2014) Ca2+-Calmodulin regulates SNARE assembly and spontaneous neurotransmitter release via v-ATPase subunit V0a1. J Cell Biol 205 (1):21–31. doi:10.1083/jcb.201312109

24. Peters C, Bayer MJ, Buhler S, Andersen JS, Mann M, Mayer A (2001) Trans-complex formation by proteolipid channels in the terminal phase of membrane fusion. Nature 409 (6820):581–588. doi:10.1038/35054500

25. Peri F, Nusslein-Volhard C (2008) Live imaging of neuronal degradation by microglia reveals a role for v0-ATPase a1 in phagosomal fusion in vivo. Cell 133 (5):916–927. doi:10.1016/j.cell.2008.04.037

26. Williamson WR, Wang D, Haberman AS, Hiesinger PR (2010) A dual function of V0-ATPase a1 provides an endolysosomal degradation mechanism in Drosophila melanogaster photoreceptors. J Cell Biol 189 (5):885–899. doi:10.1083/jcb.201003062

27. Strasser B, Iwaszkiewicz J, Michielin O, Mayer A (2011) The V-ATPase proteolipid cylinder promotes the lipid-mixing stage of SNARE-dependent fusion of yeast vacuoles. Embo J 30 (20):4126–4141. doi:10.1038/emboj.2011.335

28. Rizo J, Rosenmund C (2008) Synaptic vesicle fusion. Nature Structural & Molecular Biology 15 (7):665–674. doi:10.1038/nsmb.1450

29. Tokumaru H, Umayahara K, Pellegrini LL, Ishizuka T, Saisu H, Betz H, Augustine GJ, Abe T (2001) SNARE complex oligomerization by synaphin/complexin is essential for synaptic vesicle exocytosis. Cell 104 (3):421–432. doi:10.1016/s0092-8674(01)00229-x

30. Courtney NA, Bao H, Briguglio JS, Chapman ER (2019) Synaptotagmin 1 clamps synaptic vesicle fusion in mammalian neurons independent of complexin. Nat Commun 10 (1):4076. doi:10.1038/s41467-019-12015-w

31. Galli T, McPherson PS, De Camilli P (1996) The V0 sector of the V-ATPase, synaptobrevin, and synaptophysin are associated on synaptic vesicles in a Triton X-100-resistant, freeze-thawing sensitive, complex. J Biol Chem 271 (4):2193–2198

32. Morel N (2003) Neurotransmitter release: the dark side of the vacuolar-H+ATPase. Biology of the cell 95 (7):453–457

33. Di Giovanni J, Boudkkazi S, Mochida S, Bialowas A, Samari N, Leveque C, Youssouf F, Brechet A, Iborra C, Maulet Y, Moutot N, Debanne D, Seagar M, El Far O (2010) V-ATPase membrane sector associates with synaptobrevin to modulate neurotransmitter release. Neuron 67 (2):268–279. doi:10.1016/j.neuron.2010.06.024

34. Wang D, Hiesinger PR (2013) The vesicular ATPase: a missing link between acidification and exocytosis. J Cell Biol 203 (2):171–173. doi:10.1083/jcb.201309130

35. Rama S, Boumedine-Guignon N, Sangiardi M, Youssouf F, Maulet Y, Leveque C, Belghazi M, Seagar M, Debanne D, El Far O (2019) Chromophore-Assisted Light Inactivation of the V-ATPase V0c Subunit Inhibits Neurotransmitter Release Downstream of Synaptic Vesicle Acidification. Mol Neurobiol 56 (5):3591–3602. doi:10.1007/s12035-018-1324-1

36. David P, el Far O, Martin-Mouto N, Poupon MF, Takahashi M, Seagar MJ (1993) Expression of synaptotagmin and syntaxin associated with N-type calcium channels in small cell lung cancer. FEBS Lett 326 (1-3):135–139

37. Ramos-Miguel A, Sawada K, Jones AA, Thornton AE, Barr AM, Leurgans SE, Schneider JA, Bennett DA, Honer WG (2017) Presynaptic proteins complexin-I and complexin-II differentially influence cognitive function in early and late stages of Alzheimer’s disease. Acta neuropathologica 133 (3):395–407. doi:10.1007/s00401-016-1647-9

38. Takahashi S, Yamamoto H, Matsuda Z, Ogawa M, Yagyu K, Taniguchi T, Miyata T, Kaba H, Higuchi T, Okutani F, et al. (1995) Identification of two highly homologous presynaptic proteins distinctly localized at the dendritic and somatic synapses. FEBS Lett 368 (3):455–460

39. Nishiki T, Kamata Y, Nemoto Y, Omori A, Ito T, Takahashi M, Kozaki S (1994) Identification of protein receptor for Clostridium botulinum type B neurotoxin in rat brain synaptosomes. J Biol Chem 269 (14):10498–10503

40. Di Giovanni J, Iborra C, Maulet Y, Leveque C, El Far O, Seagar M (2010) Calcium-dependent regulation of SNARE-mediated membrane fusion by calmodulin. J Biol Chem 285 (31):23665–23675

41. Quetglas S, Leveque C, Miquelis R, Sato K, Seagar M (2000) Ca2+-dependent regulation of synaptic SNARE complex assembly via a calmodulin- and phospholipid-binding domain of synaptobrevin. Proc Natl Acad Sci U S A 97 (17):9695–9700

42. Meerbrey KL, Hu G, Kessler JD, Roarty K, Li MZ, Fang JE, Herschkowitz JI, Burrows AE, Ciccia A, Sun T, Schmitt EM, Bernardi RJ, Fu X, Bland CS, Cooper TA, Schiff R, Rosen JM, Westbrook TF, Elledge SJ (2011) The pINDUCER lentiviral toolkit for inducible RNA interference in vitro and in vivo. Proc Natl Acad Sci U S A 108 (9):3665–3670. doi:10.1073/pnas.1019736108

43. Leveque C, Boudier JA, Takahashi M, Seagar M (2000) Calcium-dependent dissociation of synaptotagmin from synaptic SNARE complexes. J Neurochem 74 (1):367–374. doi:10.1046/j.1471-4159.2000.0740367.x

44. Mochida S (1994) [Inhibition of neurotransmitter release by tetanus and botulinum neurotoxins]. Seikagaku 66 (3):254–259

45. Vitale N, Mukai H, Rouot B, Thierse D, Aunis D, Bader MF (1993) Exocytosis in chromaffin cells. Possible involvement of the heterotrimeric GTP-binding protein G(o). J Biol Chem 268 (20):14715–14723

46. Tanguy E, Coste de Bagneaux P, Kassas N, Ammar MR, Wang Q, Haeberle AM, Raherindratsara J, Fouillen L, Renard PY, Montero-Hadjadje M, Chasserot-Golaz S, Ory S, Gasman S, Bader MF, Vitale N (2020) Mono- and Poly-unsaturated Phosphatidic Acid Regulate Distinct Steps of Regulated Exocytosis in Neuroendocrine Cells. Cell Rep 32 (7):108026. doi:10.1016/j.celrep.2020.108026

47. Gabel M, Delavoie F, Demais V, Royer C, Bailly Y, Vitale N, Bader MF, Chasserot-Golaz S (2015) Annexin A2-dependent actin bundling promotes secretory granule docking to the plasma membrane and exocytosis. J Cell Biol 210 (5):785–800. doi:10.1083/jcb.201412030

48. Houy S, Estay-Ahumada C, Croise P, Calco V, Haeberle AM, Bailly Y, Billuart P, Vitale N, Bader MF, Ory S, Gasman S (2015) Oligophrenin-1 Connects Exocytotic Fusion to Compensatory Endocytosis in Neuroendocrine Cells. J Neurosci 35 (31):11045–11055. doi:10.1523/JNEUROSCI.4048-14.2015

49. Brunger AT, Leitz J (2022) The core complex of the Ca(2+)-triggered presynaptic fusion machinery. Journal of Molecular Biology:167853. doi:10.1016/j.jmb.2022.167853

50. Rama S, Boumedine-Guignon N, Sangiardi M, Youssouf F, Maulet Y, Lévêque C, Belghazi M, Seagar M, Debanne D, El Far O (2018) Chromophore-Assisted Light Inactivation of the V-ATPase V0c Subunit Inhibits Neurotransmitter Release Downstream of Synaptic Vesicle Acidification. Mol Neurobiol. doi:10.1007/s12035-018-1324-1

51. Adachi I, Puopolo K, Marquez-Sterling N, Arai H, Forgac M (1990) Dissociation, cross-linking, and glycosylation of the coated vesicle proton pump. J Biol Chem 265 (2):967–973

52. Mochida S (2011) Activity-dependent regulation of synaptic vesicle exocytosis and presynaptic short-term plasticity. Neurosci Res 70 (1):16–23. doi:https://doi.org/10.1016/j.neures.2011.03.005

53. Vitale N, Caumont AS, Chasserot-Golaz S, Du G, Wu S, Sciorra VA, Morris AJ, Frohman MA, Bader MF (2001) Phospholipase D1: a key factor for the exocytotic machinery in neuroendocrine cells. Embo J 20 (10):2424–2434. doi:10.1093/emboj/20.10.2424

54. Chasserot-Golaz S, Vitale N, Umbrecht-Jenck E, Knight D, Gerke V, Bader MF (2005) Annexin 2 promotes the formation of lipid microdomains required for calcium-regulated exocytosis of dense-core vesicles. Mol Biol Cell 16 (3):1108–1119. doi:10.1091/mbc.e04-07-0627

55. Hettie KS, Liu X, Gillis KD, Glass TE (2013) Selective catecholamine recognition with NeuroSensor 521: a fluorescent sensor for the visualization of norepinephrine in fixed and live cells. ACS Chem Neurosci 4 (6):918–923. doi:10.1021/cn300227m

56. Morel N, Dedieu JC, Philippe JM (2003) Specific sorting of the a1 isoform of the V-H+ATPase a subunit to nerve terminals where it associates with both synaptic vesicles and the presynaptic plasma membrane. J Cell Sci 116 (Pt 23):4751–4762. doi:10.1242/jcs.00791

57. Reim K, Wegmeyer H, Brandstatter JH, Xue M, Rosenmund C, Dresbach T, Hofmann K, Brose N (2005) Structurally and functionally unique complexins at retinal ribbon synapses. J Cell Biol 169 (4):669–680. doi:10.1083/jcb.200502115

58. Huntwork S, Littleton JT (2007) A complexin fusion clamp regulates spontaneous neurotransmitter release and synaptic growth. Nat Neurosci 10 (10):1235–1237. doi:10.1038/nn1980

59. Grasso EM, Terakawa MS, Lai AL, Xue Xie Y, Ramlall TF, Freed JH, Eliezer D (2022) Membrane Binding Induces Distinct Structural Signatures in the Mouse Complexin-1C-Terminal Domain. Journal of Molecular Biology:167710. doi:10.1016/j.jmb.2022.167710

60. Trimbuch T, Rosenmund C (2016) Should I stop or should I go? The role of complexin in neurotransmitter release. Nat Rev Neurosci 17 (2):118–125. doi:10.1038/nrn.2015.16

61. Chen X, Tomchick DR, Kovrigin E, Arac D, Machius M, Sudhof TC, Rizo J (2002) Three-dimensional structure of the complexin/SNARE complex. Neuron 33 (3):397–409. doi:10.1016/s0896-6273(02)00583-4

62. Reim K, Mansour M, Varoqueaux F, McMahon HT, Sudhof TC, Brose N, Rosenmund C (2001) Complexins regulate a late step in Ca2+-dependent neurotransmitter release. Cell 104 (1):71–81. doi:10.1016/s0092-8674(01)00192-1

63. Kummel D, Krishnakumar SS, Radoff DT, Li F, Giraudo CG, Pincet F, Rothman JE, Reinisch KM (2011) Complexin cross-links prefusion SNAREs into a zigzag array. Nat Struct Mol Biol 18 (8):927–933. doi:10.1038/nsmb.2101

64. Zhou Q, Zhou P, Wang AL, Wu D, Zhao M, Sudhof TC, Brunger AT (2017) The primed SNARE-complexin-synaptotagmin complex for neuronal exocytosis. Nature 548 (7668):420–425. doi:10.1038/nature23484

65. Malsam J, Seiler F, Schollmeier Y, Rusu P, Krause JM, Sollner TH (2009) The carboxy-terminal domain of complexin I stimulates liposome fusion. Proc Natl Acad Sci U S A 106 (6):2001–2006. doi:10.1073/pnas.0812813106

66. Gong J, Lai Y, Li X, Wang M, Leitz J, Hu Y, Zhang Y, Choi UB, Cipriano D, Pfuetzner RA, Sudhof TC, Yang X, Brunger AT, Diao J (2016) C-terminal domain of mammalian complexin-1 localizes to highly curved membranes. Proc Natl Acad Sci U S A 113 (47):E7590–E7599. doi:10.1073/pnas.1609917113

67. Wragg RT, Parisotto DA, Li Z, Terakawa MS, Snead D, Basu I, Weinstein H, Eliezer D, Dittman JS (2017) Evolutionary Divergence of the C-terminal Domain of Complexin Accounts for Functional Disparities between Vertebrate and Invertebrate Complexins. Front Mol Neurosci 10:146. doi:10.3389/fnmol.2017.00146

68. Snead D, Lai AL, Wragg RT, Parisotto DA, Ramlall TF, Dittman JS, Freed JH, Eliezer D (2017) Unique Structural Features of Membrane-Bound C-Terminal Domain Motifs Modulate Complexin Inhibitory Function. Front Mol Neurosci 10:154. doi:10.3389/fnmol.2017.00154

69. Mochida S (1998) Neurotoxins, cytoskeletons and calcium channels: Functional studies at mammalian synapses formed in culture. Secretory Systems and Toxins, 1st Edition edn. CRC Press,

70. Bader MF, Holz RW, Kumakura K, Vitale N (2002) Exocytosis: the chromaffin cell as a model system. Ann N Y Acad Sci 971:178–183

71. Ishizuka T, Saisu H, Odani S, Abe T (1995) Synaphin: a protein associated with the docking/fusion complex in presynaptic terminals. Biochem Biophys Res Commun 213 (3):1107–1114. doi:10.1006/bbrc.1995.2241

72. El Far O, Seagar M (2011) A role for V-ATPase subunits in synaptic vesicle fusion? J Neurochem 117 (4):603–612. doi:10.1111/j.1471-4159.2011.07234.x

73. Sharma S, Lindau M (2018) Molecular mechanism of fusion pore formation driven by the neuronal SNARE complex. Proc Natl Acad Sci U S A 115 (50):12751–12756. doi:10.1073/pnas.1816495115

74. Rothman JE, Krishnakumar SS, Grushin K, Pincet F (2017) Hypothesis - buttressed rings assemble, clamp, and release SNAREpins for synaptic transmission. FEBS Lett 591 (21):3459–3480. doi:10.1002/1873-3468.12874

